# Distinct effects of Atg3 and Atg10 on Autophagosome Formation in Yeast

**DOI:** 10.64898/2026.11.09.765410

**Authors:** Juan Salcedo, Olivia C. Clarino, Keeran Kumar, Saba Y. AlQato, S. K. Nadia Rahman Silvia, Jacquelyn R. Roberts, Mari A. Horton, Konrad Lautenschlager, Sophie Palmer, Pablo Demerath, Rakan A. Alnsour, Katherine M. Sowell, Rima S. Tout, Nick Laurent, Fadumo Hassan, Kamsiyochukwu Uwaje, Shannon Rochon, Andrea Zou, Steven K. Backues

## Abstract

Autophagosome formation involves two E2 enzymes: Atg3, which conjugates Atg8 to the lipid phosphatidylethanolamine, and Atg10, which conjugates Atg12 to Atg5. Although both are required for autophagy, it is unclear exactly what roles these pathways play in the process of autophagosome formation. We used baker’s yeast expressing mutant versions of both Atg3 and Atg10 that reduce, but do not eliminate, their activity to better understand their effects on Atg8 lipidation, autophagic flux, and the size and number of autophagosomes formed during nitrogen starvation. We found that while Atg3 primarily affects Atg8 lipidation and autophagosome size, Atg10 has less of an effect on Atg8 lipidation and primarily controls autophagosome number. This demonstrates that these two enzymes have distinct mechanistic roles in autophagosome formation, and implies an as-of-yet undiscovered function for the Atg12–Atg5 conjugate.

## Introduction

When eukaryotic cells experience nutrient limitation, they respond by upregulating pathways that break down some of their own cellular components in order to recycle those nutrients for continued survival. The best-studied of these pathways, macroautophagy (hereafter referred to simply as autophagy), involves the *de novo* creation of a membrane structure that envelops cytoplasmic cargo. This creates an autophagosome – a double-membraned vesicle with cargo inside – which then fuses with the lysosome (animals) or vacuole (other eukaryotes) to deliver that cargo for degradation and nutrient recycling^1^. Since autophagy is found throughout eukaryotes ^2,3^, and given its important role in maintaining human health^4,5^, there is great interest in understanding the molecular mechanism through which it occurs. Much of this work has utilized the model system *S. cerevisiae,* baker’s yeast, in which the first autophagy (ATG) genes were identified^6^, and where the process is best understood. Even in yeast, many questions remain about how the suite of Atg proteins initiate and guide the process of autophagosome formation and the unique membrane dynamics involved.

One of the core modules required for autophagosome formation is the Atg8 conjugation pathway. This module consists of the proteins Atg3, Atg4, Atg5, Atg7, Atg8, Atg10, Atg12, and Atg16^7,8^. The core function of this module is the attachment of Atg8, a small ubiquitin-like protein, to the autophagosome membrane by conjugating it to the lipid phosphatidylethanolamine (PE)^7^. There, Atg8 serves both in recognition of specific autophagic cargo as well as more general roles in the expansion of the phagophore - the cup-shaped membrane structure that matures into the autophagosome^7,9,10^.

The Atg8 conjugation pathway actually consists of two distinct branches that share an E1 enzyme, Atg7, but have two E2 enzymes, Atg3 and Atg10, each of which catalyzes a conjugation reaction with a separate small ubiquitin-related protein^7^ (Figure 1A) Atg10 conjugates Atg12 to Atg5, forming an isopeptide bond between the C-terminus of Atg12 and a conserved lysine in Atg5 in a process analogous to ubiquitination^11^. Atg3 conjugates Atg8 to the amine headgroup of PE in a chemically similar fashion^12^. Both Atg8 and Atg12 need to be activated by Atg7 before being handed off to their respective E2 enzymes; Atg8 additionally requires initial processing by Atg4 to expose its C-terminal glycine before it can begin this process^12^. Although there has been extensive structural and biochemical characterization of these proteins^8,13–16^, their biological role in autophagosome formation remains less clearly defined.

**Figure 1:**
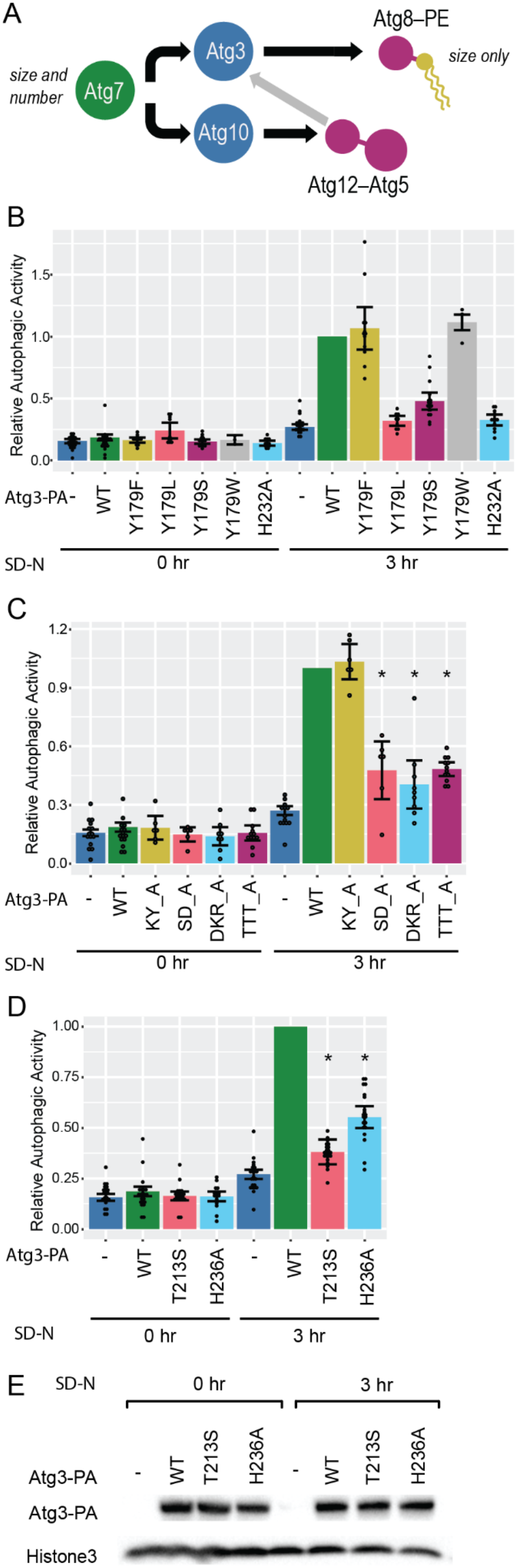
Screening for Atg3 mutants with intermediate levels of autophagy. (**A**) Simplified diagram of the Atg8–PE conjugation pathway. Atg7 activates both Atg8 and Atg12. Atg10 conjugates Atg12 to Atg5, and the Atg12–5 conjugate stimulates Atg3’s conjugation of Atg8 to PE. Surprisingly, Atg7 affects both autophagosome size and number, but Atg8 affects only autophagosome size. (**B**) Autophagic activity, as measured by the pho8Δ60 assay, caused by mutations to Y179 and H232 in the Atg3 active site. These mutations are either too strong or too weak to be useful for further study. (**C**) Autophagic activity of Atg3 double and triple mutants. KY_A: K183A, Y185A; SD_A: S206A, D208A; DKR_A: D245A, K246A, R248A; TTT_A: T21A, T24A, T25A. (**D**) Autophagic activity of Atg3 point mutants chosen for assessment of autophagosome size and number. T213S has low but significant activity, while H236A has moderate activity. (**E**) Western blot of the Protein A (PA)-tagged Atg3 wild type (WT) and mutant constructs assayed in panel C, showing that these mutants have normal expression levels. In all panels, WT or mutant Atg3-PA (with the PA tag in the flexible region) was expressed under its native promoter in an *atg3Δ* background and autophagy was assessed both in growing conditions (0 hr) and after 3 hours of nitrogen starvation in SD-N (3 hr) for induction of bulk autophagy. Error bars = 95% confidence intervals. * = significantly different (p <0.05) from both WT (1-sample T-test) and *atg3Δ* (ANOVA with Tukey HSD).

Intriguingly, both branches of this pathway appear to reconverge on Atg8–PE formation. Specifically, the Atg12–Atg5 conjugate associates with Atg16^17^, and together these activate Atg3, both helping bring it to the autophagic membrane and promoting a conformation change that stimulates Atg8–PE formation^16,18,19^. Very little Atg8–PE is formed in the absence of the Atg12–Atg5 conjugate^18,20^. This E3-like activity is the only well-established role for Atg12–Atg5 in autophagy, and is often presented as if it were the only function of this branch of the pathway^7,8,21^. However, careful analysis of the phenotypes of yeast lacking these genes does not support the hypothesis that Atg12–Atg5’s only significant role in autophagy is the stimulation of Atg8–PE formation. Some Atg8–PE, albeit much less than usual, is still formed in *atg5Δ* and *atg12Δ* cells, while no Atg8–PE formation could be detected in *atg3Δ* or *atg4Δ* cells^20^. If Atg12–Atg5’s sole role was in Atg8–PE formation, one would predict that *atg5Δ* and *atg12Δ* yeast would also have a milder defect in autophagy than *atg3Δ* or *atg4Δ* cells. But actually the opposite is true: multiple studies report a complete block in autophagy in Atg5, Atg10 or Atg12 mutants^11,22,23^, while some studies show residual autophagic activity in *atg3Δ*, *atg4Δ*, and *atg8Δ* cells^10,12,24^. Most striking is the recent data in a comprehensive re-analysis of all Atg mutants in yeast, which clearly shows only a partial block in vacuolar maturation of Ape1 (a selective autophagy substrate) in *atg3Δ*, *atg4Δ*, and *atg8Δ* cells, but a complete block in *atg5Δ*, *atg7Δ*, *atg10Δ*, and *atg12Δ* cells^25^. The same paper showed stronger autophagy defects in *atg5Δ* and *atg12Δ* than in *atg3Δ* or *atg8Δ* in a bulk autophagy assay^25^.

One strategy for gaining insight into the functions of autophagy proteins in autophagosome formation is to determine which control autophagosome size vs autophagosome number. This can be done using transmission electron microscopy (TEM) to measure the size and number of cross-sections of autophagic bodies (ABs), which serve as proxies for autophagosomes^26^. Yeast with a *pep4Δ* mutation lack protease activity in the vacuole, so the inner membrane of each autophagosome and its contents accumulate in the vacuole as ABs when bulk autophagy is stimulated by nitrogen starvation^27^. Atg8, for example, primarily controls autophagosome size. Yeast lacking Atg8 entirely form reduced numbers of very small ABs^10^, while yeast expressing lower amounts of Atg8 form the same number of ABs, but they are proportionately smaller as the amount of Atg8 is reduced^9^. Recent work has traced this effect on autophagosome size to Atg8’s ability to recruit the activating kinase, Atg1, to the surface of the expanding phagophore^28^. Similarly, it was reported that an Atg3 mutant with reduced localization to the phagophore led to fewer, smaller ABs^29^. However, smaller ABs will yield fewer cross-sections even without an actual reduction in their number, as smaller bodies are less likely to be captured in a random section, so these effects can be hard to untangle. In contrast to Atg8, cells with reduced or increased expression of Atg9 do not significantly affect AB size, but instead AB number^30^. Consistent with this, mutations in genes involved in Atg9 trafficking to the site of autophagosome formation likewise lead to the formation of fewer ABs^31–34^.

A study from our lab of Atg7, the first enzyme in the Atg8–PE conjugation pathway, showed that lower expression of Atg7 led to both smaller and fewer ABs^35^. The reduction of AB number was observed even after correcting for the sampling bias that leads to fewer sections from smaller ABs^35^. This was surprising, given that the end product of the pathway, Atg8–PE, only controls AB size, not AB number^9^. To follow up on this observation, we examined the control of autophagosome size and number by the two E2 enzymes: Atg3, which directly creates Atg8–PE, and Atg10, which creates Atg12–Atg5 and thus indirectly affects Atg8–PE formation. We hypothesized that Atg3, like Atg8, would control only autophagosome size, but that Atg10 would control both autophagosome size (via the effect of Atg12–Atg5 on Atg3 activity) as well as autophagosome number (via a novel mechanism).

## Results

### Finding Atg3 mutants with moderate reductions in autophagy

We began by tagging Atg3 with a Protein A (PA) tag to facilitate its visualization by western blot. A C-terminal PA tag blocked autophagy (Figure S1A), so we instead placed the PA in the flexible region of Atg3, based on a previous report that GFP was tolerated in this location^29^. This yielded a functional Atg3-PA construct (Figure S1B).

We then began constructing strains with altered Atg3 activity. Past size and number studies often used different promoters to control the expression level of autophagic proteins^9,30,35^; however, finding suitable promoters can be challenging, particularly for two different proteins with different expression levels. Atg3 and Atg10 are both well-studied enzymes, and many point mutations have been characterized that reduce their activity *in vitro*. Therefore, we began screening known mutants of Atg3 to find ones that led to reduced autophagic activity. We used the well-established pho8Δ60 assay, which allows a quantitative readout of the amount of cytoplasm delivered to the vacuole under starvation conditions^36^.

We were looking for mutants that led to intermediate levels of autophagic activity - less than wild type (WT) Atg3-PA, so that an effect could be seen, but more than *atg3Δ* so that sufficient ABs would be available to measure. Not all mutants that seemed promising based on their *in vitro* effects gave the desired phenotype. His232 and Tyr179 flank Atg3’s catalytic cysteine, and are conserved in Atg10, making them promising candidates^13^. However, an H232A mutation led to a complete loss of autophagic activity, as did Y179L, while Y179F and Y179W showed no loss of activity (Figure 1B). Moreover, some mutations to Tyr179 led to severely reduced Atg3 expression levels (Figure S1C), which could complicate analysis. We also tested four different double and triple mutants reported to interfere with the activation of Atg3 by Atg12–Atg5^16^. Although some of these gave intermediate levels of activity (Figure 1C), they also had some effect on expression levels (Figure S1D), and we preferred single mutations in the Atg3 active site for simplicity of interpretation. Therefore, we selected T213S and H236A for further analysis. Both Thr213 and His236 are in Atg3’s catalytic loop, and Thr213 is thought to participate directly in catalysis while His236 may help position it correctly.^14,37^ Both T213S and H236A showed levels of autophagic activity significantly lower than WT but significantly higher than *atg3Δ*; moreover, they were significantly different than each other, with H236A retaining 55% autophagic activity as compared to 38% for T213S (Figure 1D). Both mutants also retained normal Atg3-PA expression levels (Figure 1E).

### Mutations in Atg3 affect autophagosome size more than number

In order to accumulate ABs for measurement, we needed to eliminate vacuolar protease activity. Therefore, we introduced a *pep4Δ* mutation into each of these strains, and validated that this did not change Atg3-PA expression levels (Figure 2A). We tested the *in vivo* activity of Atg3 in these *pep4Δ* strains by measuring the percentage of Atg8 lipidated to Atg8–PE in both growing and starvation conditions. As expected, no Atg8–PE was detected in the absence of Atg3. The H236A mutation reduced Atg8 lipidation to only 50% (as compared to 74% in the WT), while T213S severely reduced Atg8 lipidation, to only 22% (Figure 2B-C). We then used these strains to measure the size and number of ABs formed after 3 hours of nitrogen starvation by TEM (Figure 2D). WT Atg3-PA showed large numbers of normal-sized ABs against a background of small intralumenal vesicles from the multivesicular body pathway. Some ABs were still seen in *atg3Δ* cells, but they were severely reduced in both size and number. These could be distinguished from the intralumenal vesicles both by their larger size and their smoother internal texture. The AB cross-sections from the H236A strain were, on average, significantly smaller than those seen in WT, and those in T213S were even smaller (Figure 2E). There were also significantly fewer cross-sections observed in H236A, and fewer in T213S (Figure 2F). To validate these results, we repeated this analysis on a separate set of samples generated from the same strains but starved and fixed on a different day (Figure S2A,B). This set of data was consistent with the first set; although the absolute sizes of the AB cross-sections observed was lower in all strains, they followed the same trend, with WT > H236A > T213S > *atg3Δ* (Figure 2G). Similar trends were likewise observed for the number of AB cross-sections (Figure 2H).

**Figure 2:**
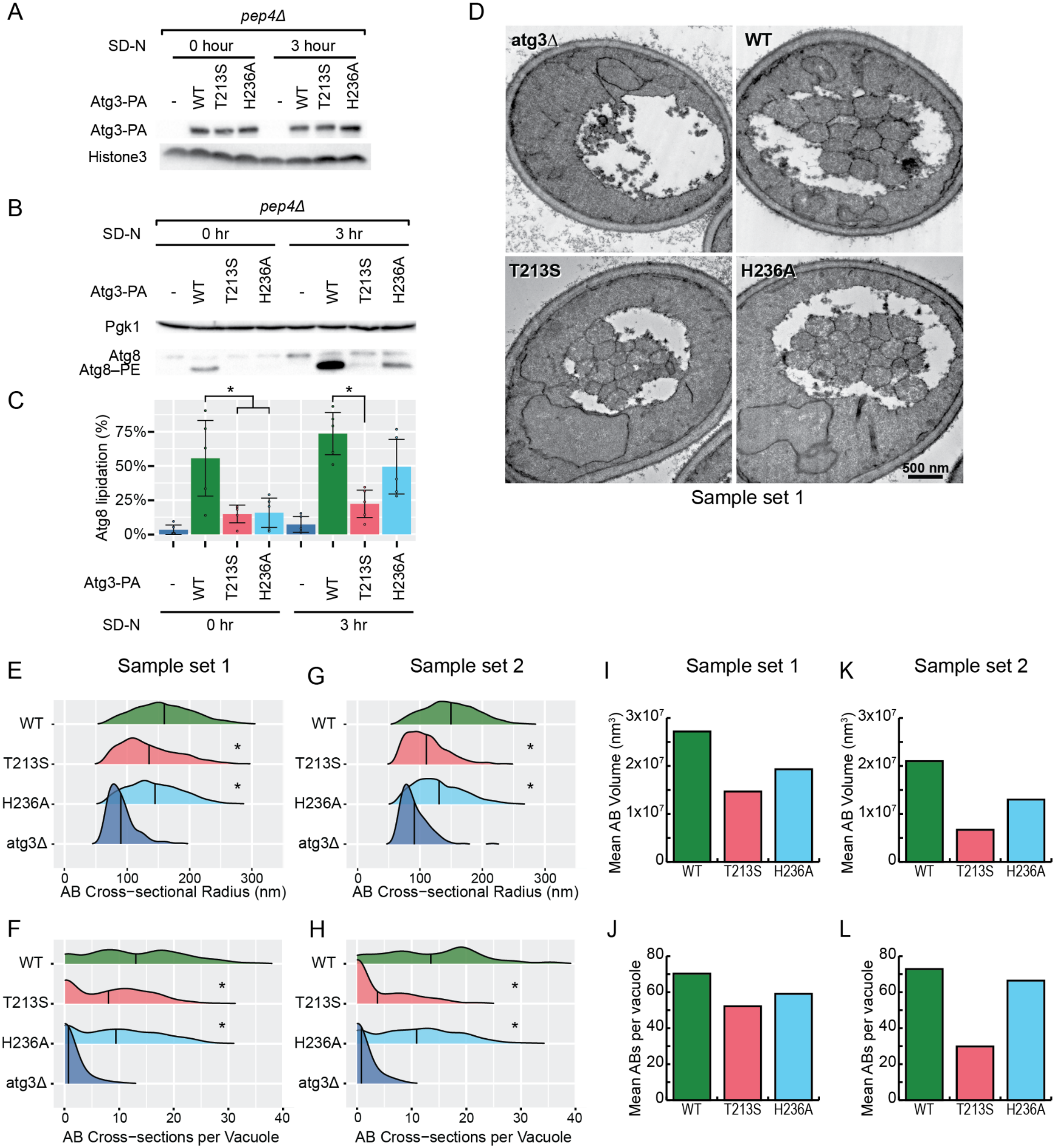
Atg3 mutants primarily affect autophagosome size. (**A**) Western blot verifying normal expression levels of Protein A (PA)-tagged Atg3 and its mutants in the four *pep4Δ* strains used for measuring autophagic bodies by electron microscopy. (**B**) Western blot showing Atg8 lipidation in these same strains in growing conditions (0 hr) and after 3 hours of nitrogen starvation in SD-N (3hr). T213S shows a severe (but not complete) defect in Atg8-PE production, while H236A has a moderate defect. (**C**) Quantification of B. Atg8-PE production is measured as % of total Atg8 that is in the lipidated Atg8-PE form. Error bars = 95% confidence intervals. * = indicated samples are significantly different (p < 0.05, ANOVA with Tukey HSD) (**D**) Representative electron micrographs of vacuole cross-sections for all four *pep4Δ* strains after 3 hours of nitrogen starvation in SD-N to induce bulk autophagy. Both mutants display smaller autophagic body cross-sections than the strain expressing WT Atg3-PA, while the atg3Δ strain contains only a few very small autophagic body cross-sections. (**E-H**) Quantification of the distribution of autophagic body cross-sectional radii (E and G) and the number of body cross-sections observed per vacuole cross section (F and H). Vertical lines in each ridgeline plot indicate the mean observed radius or body number, respectively. * = significantly different (p < 0.05) from both WT and atg3Δ; ANOVA with Tukey HSD. Data in G and H is from an independent biological replicate. N ≥ 200 images per strain per biological replicate. (**I-L**) Back-estimates of the mean volume of the autophagic bodies (I and K) and the average number per vacuole (J and L) based on the observed cross-sections. K and L represent the second biological replicate. Reliable estimates of original body size (and therefore also original body number) could not be generated for the *atg3Δ* strain because of the small number and size of body cross-sections observed.

The raw data suggested that mutations reducing Atg3’s catalytic activity reduced both autophagosome size and number. However, smaller ABs will yield fewer cross-sections even if their actual numbers are the same due to the bias of taking random slices through the vacuole. To correct for this and generate a more accurate comparison of the effect of these mutations on autophagosome size and number, we used previously developed methods^26,38^ to back-estimate the size and number of the original ABs from the cross-sectional data (Table S1). These estimates suggest that the H236A and T213S do indeed have slightly fewer ABs than WT, even when the differences in AB size are taken into account, but that the difference in AB size is much greater (Figures 2I-L). This is particularly evident when comparing AB number to mean AB volume, which is the aspect of AB size directly correlated to overall autophagic flux. We also analyzed this data using a separate method: a python-based program that slices simulated vacuoles containing spherical ABs, and compares the distributions of body cross-sections in the simulated slices to those from the real TEM data to estimate original AB size and number^39^. This analysis method yielded similar results (Figure S2C-F and Table S2). Thus, Atg3 appears to control primarily AB size, but also have a smaller effect on AB number.

### Mutations in Atg10 have a greater effect on Atg12–Atg5 conjugation than on autophagy

Next we turned to Atg10. In this case, a C-terminal PA tagged construct was functional (Figure S3A,B). The mutation of His131 to alanine caused a moderate defect in overall autophagic activity (unlike the mutation of the analogous residue, His232, in Atg3, which eliminated autophagy), while the mutation of Tyr73 (the analog to Tyr179 in Atg3) to Gln gave only a very mild defect in autophagic activity that did not reach significance (Figure S3C). Another difference between Atg3 and Atg10 is that the Y73Q mutation led to a striking overexpression of Atg10-PA (Figure S3D), which has been previously reported for various Atg10 Tyr73 mutations, though its molecular cause is unknown^15^. Both Tyr73 and His131 are conserved in human Atg10, as Tyr106 and His164, respectively (Figure S3E). Interestingly, a mutation affecting His164, H164Y, has been detected in human populations, where it is predicted to have deleterious effects^40^.

Although the Atg10 H131A mutation, with its moderate defect in autophagy, was suitable for our purposes, we also wanted a mutant with a strong (but not complete) defect in autophagy for comparison. Therefore, we combined the two active site mutants, creating a Y73Q, H131A double mutant. This retained measurable, but heavily reduced, autophagic activity via the pho8Δ60 assay (Figure 3A). To verify that there was indeed activity retained in the double mutant, we turned to the Ape1 maturation assay. This is a readout of the Cytoplasm-to-vacuole targeting (Cvt) pathway, a type of selective autophagy in yeast that delivers the precursor Ape1 (prApe1) protein to the vacuole where it is proteolytically cleaved into mature Ape1 (mApe1); this maturation can be detected by western blotting^41^. Under nitrogen starvation conditions, Ape1 is also taken up by bulk autophagosomes, though the presence of a specific Ape1 receptor means that it is taken up more efficiently than bulk cytosol. As such, it is a sensitive assay for low levels of autophagic activity. In this assay, the double mutant retained ~50% Ape1 maturation, as opposed to background levels of less than 25% in an *atg10Δ* strain, verifying that significant autophagy does still occur in the double mutant (Figures 3B,C). Thus we used the H131A single and the Y73Q, H131A double mutant for the remainder of our analysis of Atg10.

**Figure 3:**
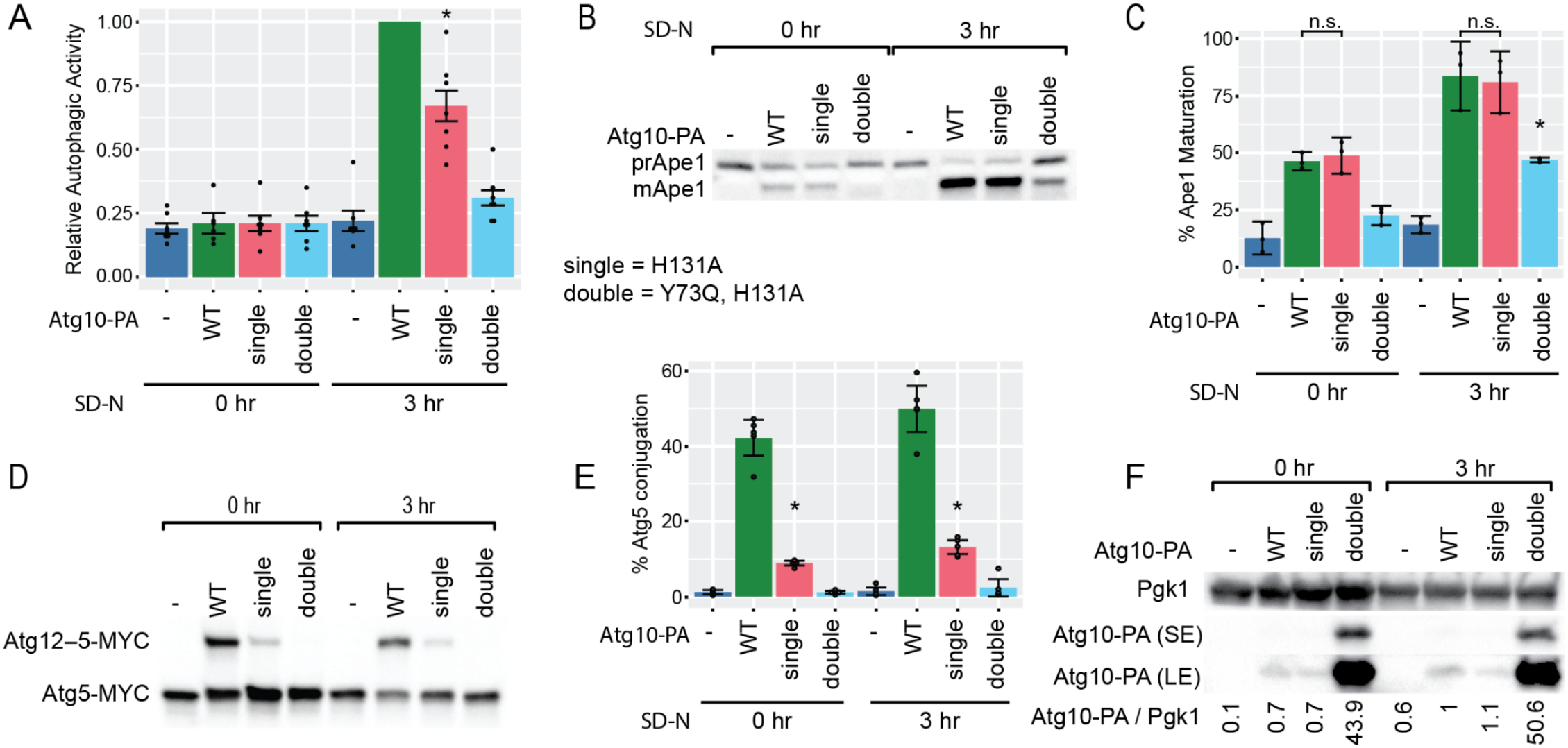
Atg10 active site mutants have a greater effect on Atg12–5 conjugation than on autophagy. **(A**) Bulk autophagic activity, as measured by the pho8Δ60 assay, in the Atg10 H131A single mutant and the Y73Q, H131A double mutant. The single mutant has a modest defect, while the double mutant shows a severe defect. * = significantly different (p <0.05) from both WT (1-sample T-test) and *atg10Δ* (ANOVA with Tukey HSD). (**B and C**) Selective autophagy (Cvt pathway), as measured by Ape1 processing. Western blotting (B) allows detection of the cytosolic form of Ape1 (prApe1) as well as the mature, vacuolar form of Ape1 (mApe1) created by the Cvt pathway; C is the quantification of % of Ape1 in its mature form from three such western blots. The H131A single mutant has no defect in the Cvt pathway, while the Y73Q, H131A double mutant has a moderate defect. (**D-E**) Quantification of Atg12–5 conjugation. Atg5-MYC is detected by western blotting against the MYC tag (D); the ATG12–5 conjugate can be seen as a higher molecular weight band. The % of Atg5 in the conjugated form in four biological replicates is quantified in E. The H131A single mutant shows a > 75% reduction in Atg12–5 conjugation, while the Y73Q, H131A double mutant has undetectable levels of the Atg12–5 conjugate. (**F**) Western blotting against the protein A (PA) tag on Atg10 shows that the Y73Q, H131A double mutant leads to much higher levels of Atg10 protein, despite the very reduced Atg10 activity level in this mutant. SE = short exposure; LE = long exposure (double mutant overexposed). Quantifications below the blot are the ratio of Atg10-PA to Pgk1, normalized to the WT 3 hr sample. In all panels, WT or mutant Atg10-PA (C-terminal tag) was integrated into the genome under its native promoter while Atg5-MYC was expressed from a centromeric plasmid in an *atg5Δ /atg10Δ* background; autophagy was assessed both in growing conditions (0 hr) and after 3 hours of nitrogen starvation in SD-N (3 hr). Error bars = 95% confidence intervals. * in panels C and E = significantly different (p <0.05) from both WT and *atg10Δ* (ANOVA with Tukey HSD)

As a direct readout of Atg10 enzymatic activity, we used western blotting to measure the amount of Atg5-MYC converted to the Atg12–Atg5 conjugate. WT Atg10-PA gave 40-50% Atg5 conjugation, while the H131A single mutant showed a four-fold reduction in activity, to only about 10% Atg5 conjugation. Atg12–Atg5-MYC levels in the double mutant were below the detection limit of our assay (Figures 3D,E). Blotting against the PA tag on Atg10 showed that the double mutant was massively overexpressed (as expected due to the mutation of Tyr73), despite having almost no enzymatic activity (Figure 3F).

### Mutations in Atg10 primarily affect autophagosome number

We introduced the *pep4Δ* mutation into each of these four strains to allow the accumulation of ABs for measurement, and verified that the expression level of Atg10-PA and its enzymatic activity as measured by Atg12–Atg5-MYC conjugation remained as expected (Figure 4A-C). To assess the effects of reducing Atg10 activity on downstream steps in the pathway, we measured Atg8 lipidation in each of these strains (Figure 4D,E). As expected, Atg8 lipidation is severely reduced, though not eliminated, in the absence of Atg10, reflecting the known role of the Atg12–Atg5 conjugate in stimulating Atg8-PE formation by Atg3. Interestingly, Atg10 single mutant shows no measurable defect in Atg8 lipidation (Figure 4D,E), despite its ~4-fold decrease in Atg12–Atg5 conjugation (Figure 4B,C). The double mutant shows a likely decrease in Atg8 lipidation after starvation (though not quite statistically significant), but is still significantly higher than that seen in *atg10Δ* (Figure 4D,E) despite the amount of Atg12–Atg5 being below detectable levels (Figure 4B,C).

**Figure 4:**
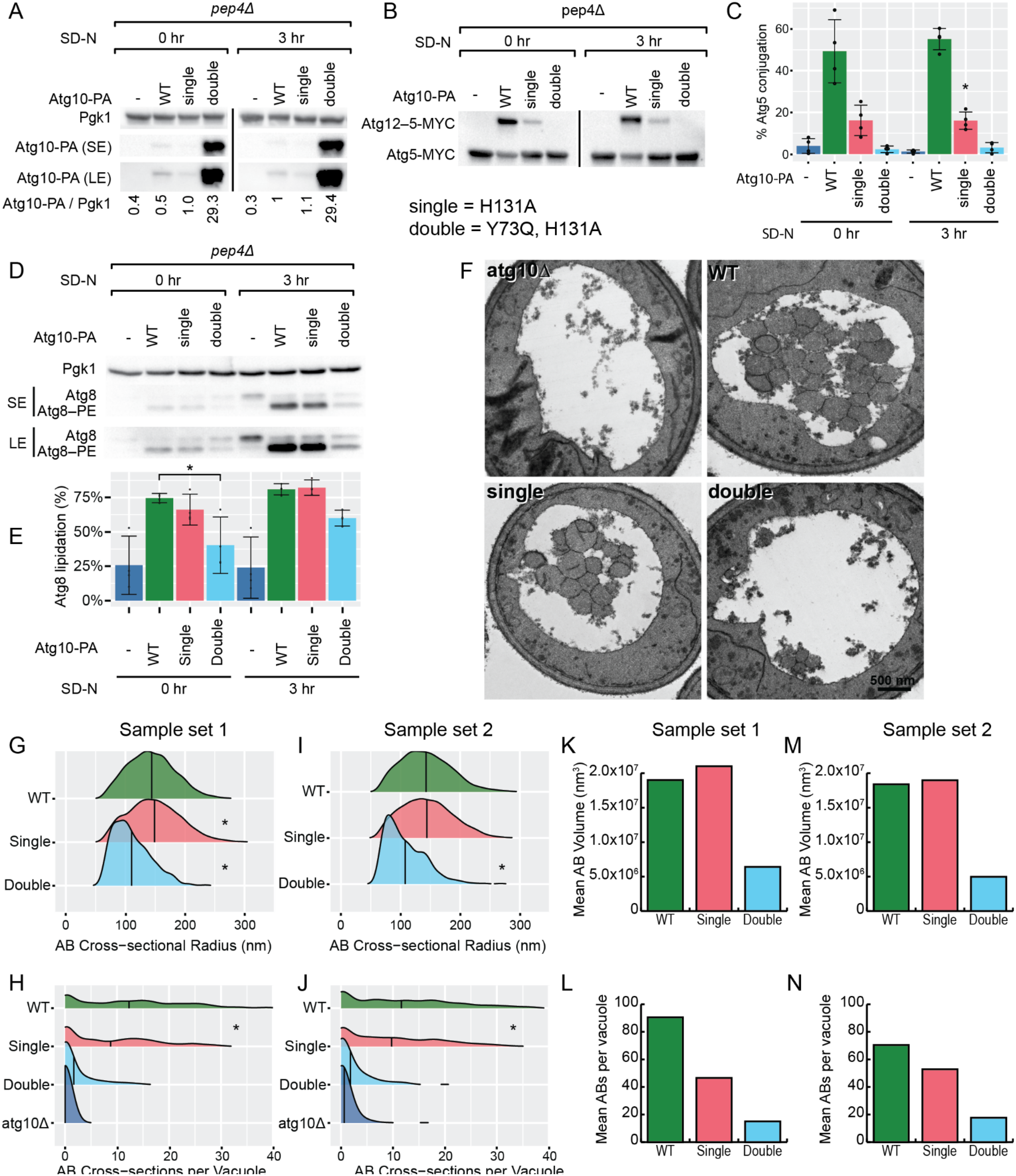
Atg10 mutants primarily affect autophagosome number. (**A**) Western blot verifying expected expression levels of Protein A (PA)-tagged Atg10 and its mutants in the *pep4Δ* strains used for measuring ABs by electron microscopy. (**B and C**) Quantification of Atg12–5 conjugation in those same strains. The % of Atg5 in the conjugated form in four biological replicates is quantified in C. The H131A single mutant shows a~70% reduction in Atg12–5 conjugation, while the Y73Q, H131A double mutant has undetectable levels of the Atg12–5 conjugate. * = significantly different (p <0.05) from both WT and atg3Δ (ANOVA with Tukey HSD). (**D**) Western blot showing Atg8 lipidation in these same strains. The H131A single mutant shows no detectable defect in Atg8 lipidation, while the Y73Q, H131A double mutant shows a moderate defect. (**E**) Quantification of B. Atg8-PE production is measured as % of total Atg8 that is in the lipidated Atg8-PE form. * = indicated samples are significantly different (p < 0.05, ANOVA with Tukey HSD) Error bars in C and E = 95% confidence intervals. (**F**) Representative electron micrographs of vacuole cross-sections for all four *pep4Δ* strains. The H131A single mutant shows fewer AB cross-sections than the WT, but those cross-sections are of similar size. The Y73Q, H131A double mutant shows far fewer and also smaller AB cross sections, while almost no AB cross-sections are seen in the *atg10Δ* strain. (**G-J**) Quantification of the distribution of AB cross-sectional radii (G and I) and the number of AB cross-sections observed per vacuole cross section (H and J). Vertical lines in each ridgeline plot indicate the mean observed radius or body number, respectively. * = significantly different (p < 0.05) from WT (G and I) or from both WT and atg3Δ (H and J); ANOVA with Tukey HSD. Data in I and J is from an independent biological replicate. N ≥ 300 images per strain per biological replicate for WT and single, and N ≥ 200 images per strain per biological replicate for double and *atg10Δ*. (**K-N**) Back-estimates of the mean volume of the ABs (K and M) and the average number per vacuole (J and L) based on the observed cross-sections. K and L represent the second biological replicate. Reliable estimates of original AB size (and therefore also original body number) could not be generated for the *atg10Δ* strain because of the extremely small number of body cross-sections observed.

We then used TEM to measure the size and number of ABs formed in these strains after 3 hours of nitrogen starvation (Figure 4F). Large numbers of normal-sized bodies were seen in the WT Atg10-PA strain. The H131A single mutant produced bodies that were slightly larger than WT, but far fewer. The double mutant, in contrast, produced much smaller bodies, and also vastly fewer (Figure 4G,H). Western blots of samples from the cultures used for TEM verified the identity of the strains (Figure S4A,B). A second sample set using the same strains but starved and fixed on a different day gave very similar results (Figure 4 I,J and Figure S4C). Back-estimates of the volume and number of the original ABs from both sample sets based on stereological formulas supported the conclusion that the single mutant had substantially fewer, slightly larger ABs than the WT, while the double mutant had even fewer, much smaller ABs (Figure 4K-N and Table S3). Analysis of the data by the python-based simulation method led to the same conclusions (Figure S4D-G and Table S4). Thus, mutations to Atg10 appear to have a primary effect on autophagosome number, as a mutation that modestly reduces Atg10 activity reduces only the number of autophagosomes formed, not their size. Only when Atg10 activity is very severely reduced does this lead to a reduction in autophagosome size along with number.

## Discussion

Our findings highlight significant differences between Atg3 and Atg10 in terms of their effect on autophagy. Reduced Atg3 activity affects both autophagosome size and autophagosome number, but the larger effect is on autophagosome size. In contrast, reduced Atg10 activity primarily affects autophagosome number; only the double mutant with very little residual Atg10 activity showed a decrease in autophagosome size. In fact, the single mutant, despite a ~35% reduction in autophagic flux as measured by the pho8Δ60 assay, actually produced slightly but significantly larger autophagic bodies than WT Atg10. Though at first sight surprising, a reciprocal relationship between autophagosome size and number upon moderate perturbation of the system has been seen before. For example, in *pho23Δ* cells, which have significantly more ABs than WT, those ABs were slightly smaller, while in an Atg9 underexpression mutant that had significantly fewer ABs than WT, those ABs were noticeably larger^30^. Similarly, both an *atg23Δ* mutant and point mutant in Atg9 that can no longer bind Atg23 give fewer but larger ABs^34^. Reduced expression of Atg8 led to smaller ABs but the same number of observed AB cross-sections, which suggests that there were actually more of the smaller ABs in order for them to be randomly sampled at the same rate^9^. This may indicate a positive correlation between the amount of time that each autophagosome takes to form and its final size, or that some other component of the autophagy pathway is limiting and so when fewer autophagosomes are made they can each become a little larger.

The Atg10 single mutant showed a four-fold reduction in the amount of Atg12–Atg5 formed, but no reduction in Atg8 lipidation. Only the Atg10 double mutant, with undetectable levels of Atg12–Atg5 present, showed less Atg8–PE, although even there it was a moderate reduction, much higher than the amount formed in an *atg10Δ* strain. This suggests that the amount of Atg12–Atg5 normally formed in the cell is well in excess of what is needed for optimal stimulation of Atg8 lipidation by Atg3.

Despite having no defect in Atg8–PE levels, the Atg10 single mutant had a ~35% reduction in autophagic activity, attributable exclusively to a reduction in autophagosome number, not autophagosome size. This is in contrast to either of the Atg3 mutants, both of which showed reductions in Atg8–PE levels as well as autophagosome size. Indeed, across all mutants tested, all of those that led to reduced Atg8–PE levels also led to smaller ABs, supporting previous reports that the amount of Atg8 directly controls autophagosome size^9,28^. The Atg3 mutants we tested also reduced autophagosome number, perhaps because of their severity, but had a larger effect on size. In contrast, the Atg10 single mutant had no effect on Atg8 lipidation and thus must have reduced autophagosome number through some alternate route. Mutation of the homolous residue in human Atg10 (H164) has been detected in human populations and predicted to have deleterious effects^40^; it would be interesting to determine if mutating this residue also decreases autophagosome number without affecting size in human cells.

Overall, this suggests that Atg10 - or, more likely, its product, the Atg12–Atg5 conjugate - has an additional, as of yet undescribed, role in autophagosome formation that specifically controls autophagosome number. What could this role be? One possibility is suggested by a 2012 report from the Martens lab that investigates the function of the Atg12–Atg5-Atg16 complex^19^. This complex has membrane-binding ability, which *in vitro* allows it not only to target Atg3 to the membrane but also to tether membranes together^19^. The *in vivo* relevance of this membrane tethering capability is not yet clear, but if it is involved in the initiation step of autophagosome formation, that would be one possible route by which Atg12–Atg5 might control autophagosome number independent from its function as an E3 for Atg8 lipidation. Another, related, possibility is suggested by a 2014 paper from the Wollert lab that found that Atg12 binds directly to Atg8 and that the Atg12–Atg5-Atg16 complex in conjunction with lipidated Atg8 forms a meshwork-like scaffold on the membrane^42^. Our Atg10 single mutant has normal levels of Atg8 lipidation but only 25% as much Atg12–Atg5, which may be too little to efficiently support this scaffold. This might plausibly lead to the reduction in autophagosome number if the scaffold is required for successful autophagosome formation, as that paper suggests^42^. However, it is surprising that this would have no effect on autophagosome size, and does not fully explain why *atg10Δ* mutants have more severe defects in autophagosome formation than *atg8Δ* mutants, since this effect is dependent on Atg8. A final possibility is suggested by a 2019 report from the Nakatogawa lab showing that the N-terminus of Atg12 interacts with the autophagy initiation complex via the scaffold protein Atg17, and that deletion of this region of Atg12 (but not other manipulations that decrease Atg8–PE formation) reduces the formation of Atg17-GFP puncta^43^. Based on this they suggested that Atg12–Atg5 may facilitate the assembly of a scaffold for the initiation of autophagosome formation, separate from its role in Atg8–PE formation^43^. Further research on Atg8-independent roles of Atg10 and the Atg12–Atg5 conjugate will be needed to distinguish between these possibilities and fully understand the functions of these proteins in autophagosome biogenesis.

## Materials and Methods

### Yeast Culture

All media components were purchased from ForMedium (Norfolk, UK) unless otherwise indicated. Peptone was from Research Products International (RPI) and glucose was from Sigma-Aldrich. All cultures were grown at 30 °C with shaking at 300 rpm.

Cultures containing Atg3-PA and its mutants were grown in YPD (yeast extract (1% w/v), peptone (2% w/v), and glucose (2% w/v)). When maintenance of the pRS414-Atg5-9xMyc plasmid was required (as in most strains containing Atg10-PA and its mutants), cultures were grown in SMD-trp (yeast nitrogen base without amino acids (0.69% w/v), glucose (2% w/v), casamino acids (0.5% w/v), adenine (0.0015% w/v), histidine (0.001% w/v), leucine (0.0025% w/v), lysine (0.0015% w/v) and uracil (0.02% w/v).

Nitrogen starvation media (SD-N) contained yeast nitrogen base without amino acids or ammonium sulfate (0.69% w/v) and glucose (2% w/v). For nitrogen starvation, log-phase cultures (OD ~ 1) grown in YPD or SMD-trp were harvested by centrifugation (5 min, 2000 x g), washed once in sterile water, and resuspended in SD-N. Starvation was performed for 3 h beginning from the onset of the water wash before harvesting the cultures.

### Plasmid and Strain Construction

All PCR reactions were carried out using Phusion HF polymerase (New England Biolabs) according to the manufacturer’s protocol in a Bio-Rad MyCycler thermal cycler. Products were resolved by agarose gel electrophoresis stained with GelRed and imaged using a Bio-Rad ChemiDoc imaging system.

All mutant plasmids were verified by Sanger sequencing (University of Michigan Genomics Core and/or Eton Biosciences). Yeast transformations were performed via the LiAc/SS carrier DNA/PEG transformation method^44^. Primers used are listed in Table 1, plasmids in Table 2, and strains in Table 3

**Table 1:** Plasmids Used in this Study.

| Name | Description | Source |
|---|---|---|
| pRS406 | Empty vector | Ref<sup>55</sup> |
| pRS406-Atg3-PA-FR | Native-promoter driven ATG3 with PA tag between residues 117 and 118 | This study |
| pRS406-Atg3-PA-FR<sup>Y179F</sup> | Atg3 single mutant | This study |
| pRS406-Atg3-PA-FR<sup>Y179L</sup> | Atg3 single mutant | This study |
| pRS406-Atg3-PA-FR<sup>Y179S</sup> | Atg3 single mutant | This study |
| pRS406-Atg3-PA-FR<sup>Y179W</sup> | Atg3 single mutant | This study |
| pRS406-Atg3-PA-FR<sup>T213S</sup> | Atg3 single mutant | This study |
| pRS406-Atg3-PA-FR<sup>H236A</sup> | Atg3 single mutant | This study |
| pRS406-Atg3-PA-FR<sup>H232A</sup> | Atg3 single mutant | This study |
| pRS406-Atg3-PA-FR-TTT_A | Atg3 triple mutant (T21A, T24A, T25A) | This study |
| pRS406-Atg3-PA-FR-DKR_A | Atg3 triple mutant (D245A, K246A, R248A) | This study |
| pRS406-Atg3-PA-FR-KY_A | Atg3 double mutant (K183A, Y185A) | This study |
| pRS406-Atg3-PA-FR-SD_A | Atg3 double mutant (S206A, D208A) | This study |
| pRS406-Atg10-PA | Native-promoter-driven Atg10 with a C-terminal PA tag | This study |
| pRS406-Atg10-PA<sup>H131A</sup> | Atg10 single mutant | This study |
| pRS406-Atg10-PA<sup>Y73Q</sup> | Atg10 single mutant | This study |
| pRS406-Atg10-PA<sup>Y73Q, H131A</sup> | Atg10 double mutant | This study |
| pRS416-Atg5pro-Atg5-9xMyc-cyc1term | Native-promoter-driven Atg5 with 9xMyc tag | Ref<sup>19</sup> |
| pRS414-Atg5pro-Atg5-9xMyc-cyc1term | Native-promoter-driven Atg5 with 9xMyc tag | This study |

**Table 2:** Strains Used in this Study.

| Name | Genotype | Source |
|---|---|---|
| SEY6210 | MATα *his3Δ200 leu2-3,112 lys2-801 suc2-Δ9 trp1Δ901 ura3-52* | Ref<sup>56</sup> |
| SKB680 | WLY176 *Atg10-PA::LEU2* | This study |
| SKB890 | WLY176 *atg3Δ::TRP1* | This study |
| SKB1008 | WLY176 *atg10Δ::KANMX* | This study |
| SKB1036 | WLY176 *atg3Δ::TRP1 ura3-52::pRS406-ATG3-PA-FR* | This study |
| SKB1038 | WLY176 *atg3Δ::TRP1 ura3-52::pRS406-ATG3<sup>H232A</sup>-PA-FR* | This study |
| SKB1040 | WLY176 *atg3Δ::TRP1 ura3-52::pRS406* | This study |
| SKB1042 | WLY176 *atg3Δ::TRP1 ura3-52::pRS406-ATG3<sup>Y179F</sup>-PA-FR* | This study |
| SKB1044 | WLY176 *atg3Δ::TRP1 ura3-52::pRS406-ATG3<sup>Y179S</sup>-PA-FR* | This study |
| SKB1078 | WLY176 *atg10Δ::KANMX atg5Δ ura3-52::pRS406* | This study |
| SKB1080 | WLY176 *atg10Δ::KANMX atg5Δ ura3-52::pRS406-ATG10-PA* | This study |
| SKB1081 | WLY176 *atg10Δ::KANMX atg5Δ ura3-52::pRS406-ATG10<sup>H131A</sup>-PA-FR* | This study |
| SKB1083 | WLY176 *atg10Δ::KANMX atg5Δ ura3-52::pRS406-ATG10<sup>Y73Q, H131A</sup>-PA* | This study |
| SKB1100 | WLY176 *atg3Δ::TRP1 ura3-52::pRS406-ATG3<sup>Y179L</sup>-PA-FR* | This study |
| SKB1102 | WLY176 *atg3Δ::TRP1 ura3-52::pRS406-ATG3<sup>Y179W</sup>-PA-FR* | This study |
| SKB1128 | WLY176 *atg10Δ::KANMX atg5Δ pep4Δ::LEU2 ura3-52::pRS406* | This study |
| SKB1129 | WLY176 *atg10Δ::KANMX atg5Δ pep4Δ::LEU2 ura3-52::pRS406-ATG10-PA* | This study |
| SKB1130 | WLY176 *atg10Δ::KANMX atg5Δ pep4Δ::LEU2 ura3-52::pRS406-ATG10<sup>H131A</sup>-PA* | This study |
| SKB1131 | WLY176 *atg10Δ::KANMX atg5Δ pep4Δ::LEU2 ura3-52::pRS406-ATG10<sup>Y73Q/H131A</sup>-PA-FR* | This study |
| SKB1142 | WLY176 *atg3Δ::TRP1 ura3-52::pRS406-ATG3<sup>H236A</sup>-PA-FR* | This study |
| SKB1144 | WLY176 *atg3Δ::TRP1 ura3-52::pRS406-ATG3<sup>T213S</sup>-PA-FR* | This study |
| SKB1177 | WLY176 *pep4Δ::LEU2 atg3Δ::TRP1 ura3-52::pRS406* | This study |
| SKB1178 | WLY176 *atg3Δ::TRP1 pep4Δ::LEU2 ura3-52::pRS406-ATG3-PA-FR* | This study |
| SKB1179 | WLY176 *atg3Δ::TRP1 pep4Δ::LEU2 ura3-52::pRS406-ATG3<sup>T213S</sup> -PA-FR* | This study |
| SKB1180 | WLY176 *atg3Δ::TRP1 pep4Δ::LEU2 ura3-52::pRS406-ATG3<sup>H236A</sup> -PA-FR* | This study |
| SKB1182 | WLY176 *atg3Δ::TRP1 pep4Δ::LEU2 ura3-52::pRS406-ATG3<sup>D245A,K246A,R248A</sup>-PA-FR* | This study |
| SKB1184 | WLY176 *pep4Δ::LEU2 atg3Δ::TRP1 ura3-52::pRS406-ATG3<sup>T21A,T24A,T25A</sup>-PA-FR* | This study |
| SKB1201 | WLY176 *atg3Δ::TRP1 pRS406-ATG3<sup>K183A,Y185A</sup>-PA-FR* | This study |
| SKB1204 | WLY176 *atg3Δ::TRP1 pRS406-ATG3<sup>S206A,D208A</sup>-PA-FR* | This study |
| WLY176 | SEY6210 *pho13Δ pho8Δ60::HIS3* | Ref<sup>57</sup> |

**Table 3:** Antibodies Used in this Study.

| **Name** | **Identifier (Source)** | **Concentration** |
|---|---|---|
| Peroxidase anti-peroxidase (PAP) | Jackson ImmunoResearch, 323-005-024 | 1:5,000 |
| rabbit-α-Histone H3 | Abcam ab1791 | 1:1,000 |
| mouse-α-Pgk1 | Abcam, ab113687 | 1:10,000 |
| mouse-α-Dpm1 | Invitrogen, A-6429 | 1:1,000 |
| mouse-α-MYC | MilliporeSigma, 05-724 | 1:1,000 |
| rabbit-α-Ape1 | Daniel Klionsky | 1:5,000 |
| mouse-α-Atg8(Aut7) | Santa Cruz Biotechnology, sc-373963 | 1:500 |
| goat-α-rabbit 2˚HRP | Jackson ImmunoResearch, 111-035-003 | 1:5,000 |
| goat-α-mouse 2˚HRP | Jackson ImmunoResearch, 115-035-003 | 1:5,000 |

#### pRS414-Atg5-MYC Plasmid construction

To enable detection of Atg12–Atg5 conjugates by immunoblotting, a TRP1-marked plasmid expressing C-terminally 9xMyc-tagged Atg5 was generated. The ATG5 promoter, ATG5 coding sequence, 9xMYC tag, and CYC1 terminator were amplified (using primers SKB547 and SKB548) from pRS316-Atg5pro-Atg5-9xMyc-cyc1term obtained from the Martens lab^19^. The pRS414 vector was digested with EcoRI and BamHI, and the Atg5-9xMyc insert was cloned into the digested vector by In-Fusion cloning to generate pRS414-Atg5pro-Atg5-9xMyc-cyc1term. The In-Fusion product was transformed into *E. coli*, transformants were selected on LB-carbenicillin plates, and candidate plasmids were verified by Sanger sequencing using primers SKB19 and SKB20 before use in yeast transformation.

#### Strain construction for ATG3

A series of yeast strains were constructed in the WLY176 background to allow for quantitative measurement of autophagic activity using the pho8Δ60 assay. First, the *ATG3* gene was replaced with a *TRP1* selection marker by homologous recombination using primers SKB400 and SKB401 and template plasmid pFA6-trp^45^. Transformants were selected on SMD-trp plates, yielding the *atg3Δ* strain SKB890.

To enable detection of Atg3 by immunoblotting, an internal Staphylococcal Protein A (PA) tag was inserted into the flexible region of Atg3 between residues Gln117 and Ser118. ATG3, including 1 kb of flanking sequence, was amplified using primers SKB421 and SKB422 and cloned into pRS406, linearized with EcoRI and HindII, using In-Fusion Eco Dry Cloning to generate pRS406-ATG3pATG3. This plasmid was linearized by PCR using primers SKB425 and SKB426, and a PA tag amplified from pRS416-pCU-PA^46^ using primers SKB427 and SKB441 was inserted by In-Fusion to generate pRS406-ATG3-PA-FR. The plasmid was linearized and integrated into the *ura3-52* locus of the *atg3Δ* strain (SKB890), transformants were selected on SMD-ura plates, and PA expression was confirmed by immunoblotting with PAP antibody. This construct and an empty pRS406 vector control were integrated in parallel, generating strains SKB1036 (WT) and SKB1040 (empty).

Point mutants Y179F, Y179S, and H232A were generated by site-directed mutagenesis of pRS406-ATG3-PA-FR using primer pairs SKB492/SKB493, SKB494/SKB495, and SKB477/SKB478, respectively. PCR products were circularized via In-Fusion Eco Dry Cloning. Additional point, double, and triple mutants were synthesized in the pRS406-ATG3-PA-FR plasmid by GenScript Biotech. These included point mutants T213S, H236A, Y179L, and Y179W; double mutants KY_A (K183A,Y185A) and SD_A (S206A,D208A); and triple mutants TTT_A (T21A,T24A,T25A) and DKR_A (D245A,K246A,R248A). All plasmids were integrated into the *ura3-52* locus of SKB890.

To enable TEM imaging of intact autophagic bodies, *pep4Δ::LEU2* was introduced into strains SKB1040 (empty), SKB1036 (WT), SKB1144 (T213S), and SKB1142 (H236A) by transforming a SacI-linearized pep4Δ::LEU2 plasmid (Klionsky Lab). Transformants were selected on SMD-leu plates.

#### Strain Construction for ATG10

First, the *ATG10* gene was replaced with a kanamycin resistance cassette by homologous recombination using primers SKB399 and SKB434 and template plasmid pFA6-KanMX6^45^. Transformants were selected on YPD plates supplemented with G418. This yielded strain SKB1000 (WLY176 *atg10Δ::kan*).

To enable detection of Atg10 by immunoblotting, a C-terminal Staphylococcal protein A (PA) tag was introduced. Atg10 was first genomically tagged with PA using plasmid pFA6-PA-KanMX6 and primers SKB398 and SKB399. Atg10-PA, driven by the native *ATG10* promoter, was then amplified using primers SKB423 and SKB309 and cloned into pRS406, linearized with EcoRI and HindII, using In-Fusion Eco Dry Cloning to generate pRS406-Atg10-PA. The plasmid was linearized with StuI within the *URA3* gene and integrated into the *ura3-52* locus of SKB1000. Transformants were selected on SMD-ura plates, and PA tag expression was confirmed by immunoblotting with PAP antibody.

Single point mutants were generated by site-directed mutagenesis of pRS406-Atg10-PA. The Y73Q mutation was introduced using primers SKB490 and SKB491, generating pRS406-Atg10-PA Y73Q, and the H131A mutation was introduced using primers SKB488 and SKB489, generating pRS406-Atg10-PA H131A. A double mutant, pRS406-Atg10-PA H131A/Y73Q, was generated by introducing the H131A mutation into pRS406-Atg10-PA Y73Q using the same primers. For all mutations, the PCR product was treated with DpnI at 37°C to destroy the original template, purified, and circularized using In-Fusion Eco Dry Cloning.

To enable TEM imaging of intact autophagic bodies, *pep4Δ::LEU2* was introduced into strains SKB1078 (empty), SKB1080 (WT), SKB1081 (H131A), and SKB1083 (H131A/Y73Q) by transforming a SacI-linearized pep4Δ::LEU2 plasmid. Transformants were selected on SMD-leu plates.

#### PCR-based genotyping

PCR-based genotyping was performed on genomic DNA isolated from liquid cultures^47^ of all transformants to confirm successful strain construction.

Deletion of *ATG3* was verified using two independent primer pairs. Primers SKB429 and SKB426 confirmed the absence of the wild-type *ATG3* locus. The presence of the pFA6A-derived *TRP1* disruption cassette inserted in place of *ATG3* was detected using primers SKB429 and SKB444. Transformants producing a band with SKB429/SKB444 but no band with SKB429/SKB426 were confirmed as *atg3Δ::trp*.

Deletion of *ATG10* was verified in a similar manner. Primers SKB481 and SKB482 confirmed the absence of the wild-type *ATG10* locus. The presence of the kanamycin resistance cassette inserted in place of *ATG10* was detected using primers SKB444 and SKB482. Transformants producing a band with SKB444/SKB482 but no band with SKB481/SKB482 were confirmed as *atg10Δ::kan*.

Similar methods were used for the verification of the deletion of *PEP4*. The absence of *PEP4* was verified using primers SKB350 and SKB352, while the presence of the *LEU2* selection marker inserted in place of *PEP4* was confirmed using primers SKB350 and SKB351. Confirmed transformants were re-struck four times on fresh selective plates to ensure complete loss of residual protease activity, which was subsequently verified by immunoblotting for Ape1 processing.

### Pho8Δ60 Assays

Bulk autophagy was quantified using the Pho8Δ60 alkaline phosphatase assay following a previously established protocol^36^ with the minor modification that during sample preparation, PMSF was excluded to prevent interference with protein quantification. Protein concentrations were determined using a bicinchoninic acid (BCA) assay (ThermoFisher) carried out in a 96-well microplate format, and absorbance was measured using a BioTek Synergy 2 plate reader. Pho8Δ60 enzymatic activity was then normalized to total protein levels to account for differences in cell lysis efficiency and sample loading.

### Western Blotting

Yeast samples for western blotting were taken either in growing conditions or after 3 hours of starvation in SD-N. One OD of culture was precipitated with ice-cold 10% TCA followed by washing with ice-cold acetone. Pellets were dissolved with vortexing on a disrupter genie with 0.5mm glass beads in 50μl 1x SSB supplemented with 0.25M Tris-HCl pH 6.8, heated 95℃ for 5 minutes, centrifuged for 5 min, and then loaded (10 µl per lane) on a Mini-PROTEAN 4-15% precast polyacrylamide gel (Bio-Rad, 4561086). Separation and transfer were performed with Bio-Rad SDS-PAGE and tank transfer apparatuses following standard procedures. Proteins were transferred to Immobilon-P PVDF membranes (0.45 µm; Millipore, IPVH00010) using a Towbin-EtOH transfer buffer (15 mM Tris, 192 mM glycine, 9.5% v/v ethanol). BLUEStain 2 protein ladder (Goldbio) was used for molecular mass estimation.

All blocking and antibody incubations were performed in blocking buffer: 4% w/v nonfat dry milk prepared in TBS-T (20mM Tris, 150mM NaCl, 0.1% v/v Tween-20, pH 6.6). Membranes were blocked for 1 hour at room temperature, then incubated overnight at 4°C with gentle shaking in primary antibody diluted in blocking buffer. Antibodies and dilutions are listed in Table 3. Following primary antibody incubation, membranes were washed four times for 5 minutes each in TBS-T with gentle shaking. All secondary antibody incubations were performed for 1 hour at room temperature with gentle shaking, followed by washing four times for 5 minutes each in TBS-T with gentle shaking. Detection was performed using either Immobilon Crescendo Western HRP substrate (MilliporeSigma, WBLUR0100) or SuperSignal West Pico PLUS Chemiluminescent Substrate (Thermo Scientific, 34577), and imaged using a Bio-Rad ChemiDoc imaging system.

In some cases, blots initially probed with mouse-α-MYC antibodies were subsequently stripped by incubation in Tris-EDTA stripping buffer (0.1 M Tris, 0.01 M EDTA, 0.01% v/v Tween-20, pH 9.2) at 95℃ for 10 minutes following a published protocol^48^, then reprobed with rabbit-α-Ape1 antibodies to measure Ape1 processing.

Atg8 and Atg8-PE immunoblotting was performed as described above, except that samples were resolved on hand-poured urea SDS-PAGE gels to improve separation of free Atg8 from Atg8–PE. These gels consisted of a 13% resolving gel with 36% w/v urea and a 4% stacking gel. Gels were prepared using 30% acrylamide/bis-acrylamide 37.5:1 (Bio-Rad, 1610158), resolving gel buffer for PAGE, pH8.8 (Bio-Rad, 1610798), stacking gel buffer for PAGE, pH 6.8 (Bio-Rad, 1610799), urea (Bio-Rad, 1610730), SDS (Bio-Rad, 1610301), ammonium persulfate (Bio-Rad, 1610700), and TEMED (Bio-Rad, 1610800).

Quantification of western blots was performed on the original .scn files with ImageJ 1.54p^49^, FIJI distribution^50^, using only bands in the linear range (visible but non-saturated, as determined by the ImageLab software provided with the BioRad Geldoc).

### TEM imaging

Transmission electron microscopy (TEM) sample preparation was performed largely as previously described^26^, with minor modifications to improve membrane preservation. Briefly, log-phase yeast cultures subjected to 3 hours nitrogen starvation were fixed with potassium permanganate (KMnO₄). Cell pellets from 30 ODs of cells were resuspended in 2 mL KMnO₄ in a 50 mL conical tube and then divided equally into two microcentrifuge tubes and incubated at room temperature for 1 h. Fresh 1.5% KMnO₄ was added and samples were incubated at 4 °C for 1 h. The fixative was then replaced with fresh 1.5% KMnO₄ and samples were incubated overnight at 4 °C. Following fixation, the pellets were recombined into a single tube prior to dehydration in acetone, embedding in Spurr’s resin, and ultrathin sectioning (70 nm). Imaging was performed on a JEOL JEM 1400PLUS (LaB6) TEM at 80 keV using an AMT NanoSprint 12 scientific grade CMOS camera. Individual cells for AB labeling were captured at 6000x, while images for labeling of entire vacuoles were captured at 800x. A minimum of 199 individual cell images for AB labeling and 16 images for vacuole labeling were analyzed for each strain.

Vacuoles, including any internal ABs, were manually outlined in Photoshop or GIMP and measured in ImageJ (FIJI) as described^26^.

### Measurement of ABs with a custom cellpose model

Cellpose2 is a published machine learning computer vision platform^51^ for which we have trained a model that can efficiently label individual ABs in TEM images^52^. Raw TEM images were resized from 4112 x 3788 pixels to 2240 x 2504 pixels so the size of autophagic bodies in pixels matched the model’s training data. Labeling was performed using the “Cellpose Autophagy Body model t_2_20” running in the Cellpose2.1.1_cv2 Biocontainer in Docker via previously published^52^ Python code (https://github.com/StevenBackues/cellpose_AB) running in Pycharm. All masks generated by the model were reviewed by at least one human scientist for accuracy. The masks were measured to generate comma separated values files of the area of each cross section and the number of cross sections per imaged vacuole via previously published Python code^52^.

### Estimate of original of AB Size and Number via Stereological Calculations

Measurements of the validated masks of the AB cross-sections were used to estimate the size distribution of the original ABs using stereological formulas implemented in Microsoft Excel as previously described^26^. These calculations model the ABs as perfect spheres whose radii follow a log-normal distribution defined by mu and sigma. For this analysis, the AB cross-section recognition threshold was set to “Ignore Below” = 65nm, “Full Recognition Above” = 75 nm, as this helped to reduce interference from the multivesicular bodies present in the vacuoles of these strains. Expected distributions of cross-sectional radii were calculated for various values of mu and sigma, and then a linear regression was performed of mu and sigma against the mean and standard deviation of the experimentally observed cross-sectional radii. This yielded a best-fit mu and sigma for the original ABs, which was then used to calculate their average volume.

The same methods were used to estimate the size distribution of the original vacuoles from measurements of vacuole outlines. The vacuole cross-section recognition thresholds were set to “Ignore Below” = 100nm, “Full Recognition Above” = 250 nm for the Atg10 set 1 samples and “Ignore Below” = 300nm, “Full Recognition Above” = 350 nm for the Atg10 set 2 samples and all of the Atg3 samples to best match the conventions used while imaging.

Original AB number was estimated based on the observed number of AB cross-sections per vacuole cross-section, taking into account the AB size distribution and the average vacuole radius estimated above. This was implemented in Microsoft Excel as previously described^26^.

### Graphing and statistical analysis

Graphs were generated in R (version 4.5.1) using the ggplot2 library^53^. Statistical analysis was performed in Microsoft Excel. Error bars are 95% confidence intervals and were calculated using the “CONFIDENCE.T” function. One-sample T-tests were performed by calculating the t-statistic as (100%-experimental average) / Standard Error of the Mean, then using the 2-tailed TDIST function to find the p value. One-factor ANOVA with Tukey’s HSD post-hoc test was performed in Microsoft Excel using the Real Statistics Resource Pack software (Release 9.8). Copyright (2013 – 2026)^54^.

## Abbreviations

AB: Autophagic Body
Cvt: Cytoplasm-to-vacuole targeting
PA: Protein A
PE: phosphatidylethanolamine
TEM: Transmission Electron Microscopy
WT: Wild-type

## Acknowledgements

Thanks to Dr. Daniel Klionsky at the University of Michigan for helpful advice and rabbit-⍺-Ape1 antibodies.

Thanks to Dr. Sascha Martens at the University of Vienna for the kind gift of the pRS416-Atg5pro-Atg5-9xMyc-cyc1term plasmid. Thanks to the University of Michigan Medical School Microscopy Core for use of their TEM and advice on sample preparation.

## Funding

Supported by National Science Foundation RUI grant # 2243163 to Steven K. Backues

## Disclosure Statement

No potential conflict of interest was reported by the authors.

## Notes

### Competing Interest Statement

The authors have declared no competing interest.

