## Supplemental Methods, Tables, and Figures for "Distinct effects of Atg3 and Atg10 on Autophagosome Formation in Yeast"

***Supplemental Data for Salcedo et al. "Distinct effects of Atg3 and Atg10 on Autophagosome Formation in Yeast"***

### Supplemental Methods

**Estimate of original of AB Size and Number via a python-based simulation**

To help validate the estimates obtained from the stereological formulas, we developed a pixel-based python simulation called Autophagic Vacuole Simulator (AVS) that generates spherical ABs within a spherical vacuole, using a log-normal distribution for both AB size and number as well as vacuole size, then takes a virtual slice and calculates the radius and number of the observed AB cross-sections captured in the slice. Fitting the distribution of the real cross-sectional measurements to those of the virtual cross-sections generated with various parameters allows the estimation of the original size and number of the ABs. A full description of this approach and its validation was recently published (REF). Here, we used this simulation as an alternate approach to estimate the size and number of the ABs from each of our conditions to ensure that both analysis methods still yielded the same qualitative results.

We began by running AVS in "size mode" to generate estimated mu and sigma values for the AB size distribution from each sample set. 10,000 virtual vacuoles were generated and sliced for each combination of mu and sigma. To determine the "best fit" mu and sigma for each set of real data, we used a combination of 1) linear regression of mu and sigma against mean and standard deviation of the observed slice radii 2) Minimization of two different statistics of best fit comparing the observed and simulated slice radii distributions - the Kolmogorov Smirnov statistic and the Epps-Singleton statistic, and 3) visual inspection of QQ-plots and cumulative density function plots. Simulation runs with different values of mu and sigma were systematically tested in increments of 0.1 to ensure that all estimates fell within the range of the values tested.

Then, we ran AVS in number mode (varying the number of ABs per vacuole while holding the AB size distribution constant) to estimate original AB number. For these runs, we used the AB size mu and sigma values calculated for each sample set from running the simulation in size mode, and the vacuole size mu and sigma vacuoles derived from the stereological calculation methods. Here, AB number sigma was fixed at 0.4 to save computation time, while AB number mu was allowed to vary over a range of at least 0.9, in increments of 0.1. 10,000 virtual vacuoles were generated and sliced for each mu value. The final value of AB number mu that we used to estimate the number of ABs per vacuole was a simple average of 1) the estimates given by linear regression of mu vs mean observed body number, 2) the mu value of the run that gave the lowest Kolmogorov Smirnov statistic, and 3) the mu value of the run that gave the lowest Epps-Singleton statistic.

AVS was run using python 3.12 on custom Amazon Web Services servers, as described (Ref readme of Github). We used option "4" in the AVS main interface to simulate the bodies as perfect spheres. The following parameters were used for all simulation runs:

- Sample_size: 10,000 virtual vacuoles per mu and sigma combination
- Scale_Factor: 8 nm per voxel side
- pvals: 2, to give spherical bodies
- Optimmaxiter: 0 (no extra clustering)
- Show_Wall: False (vacuole wall not explicitly modeled in the simulation)
- unScaledSliceThickness: 70 nm (the thickness of the ultrathin sections used for TEM)
- unScaledVacMin (vacuole recognition limit): 100 nm for the Atg10 set 1 samples and 300 nm for the Atg10 set 2 samples and all of the Atg3 samples
- unScaledminBodyRadius (AB recognition limit): 65 nm
- sliceLocation: predetermined

**Sequence Alignment**

Sequence alignment was performed with Clustal Omega version 1.2.4 (https://www.ebi.ac.uk/jdispatcher/msa/clustalo)[1]. The yeast Atg10 sequence were sourced from the Saccharomyces Genome Database, SGD (https://www.yeastgenome.org/locus/S000003965#sequence), and the human Atg10 sequence was from UniProt (https://rest.uniprot.org/uniprotkb/Q9H0Y0.fasta)


**Table S1: Observed and estimated parameters from Atg3 TEM data**

Estimates are based on stereological calculations, as described in Backues et al. 2014[26]

| | | | atg3Δ | | ATG3 WT | |
|---|---|---|---|---|---|---|
| | | | **Set 1** | **Set 2** | **Set 1** | **Set 2** |
| AB Size | Observed AB cross-sectional radii | Mean (nm) | 90.1 | 91.0 | 159.7 | 149.7 |
| | | st. dev. (nm) | 21.2 | 21.5 | 47.8 | 42.3 |
| | Estimated original lognormal distribution of AB sizes | mu | N/A | N/A | 5.15 | 5.07 |
| | | sigma | N/A | N/A | 0.236 | 0.224 |
| | | mean original radius (nm) | N/A | N/A | 176.4 | 163.6 |
| | | st. dev. of orig. radii (nm) | N/A | N/A | 42.2 | 37.1 |
| | | mean orig. volume ($nm^3$) | N/A | N/A | $2.7x10^7$ | $2.1x10^7$ |
| Vacuole Size | Observed vacuole cross-sectional radii | mean (nm) | 736.9 | 766.5 | 920.4 | 866.3 |
| | | st. dev. (nm) | 223.5 | 211.7 | 280.8 | 242.6 |
| | | sample size | 519 | 298 | 162 | 314 |
| | Estimated original lognormal distribution of vacuole sizes | mu | 6.69 | 6.76 | 6.97 | 6.90 |
| | | sigma | 0.243 | 0.192 | 0.233 | 0.191 |
| | | mean original radius (nm) | 833 | 875 | 1092 | 1010 |
| ABs per vacuole | Observed AB cross-sections per vacuole | total AB cross-sections | 166 | 185 | 2661 | 2823 |
| | | vacuoles counted | 236 | 230 | 204 | 207 |
| | | observed ABs / vacuole | 0.70 | 0.80 | 13.0 | 13.6 |
| | Estimated original number of ABs per vacuole | | N/A | N/A | 70.3 | 72.8 |

| | | | ATG3 T213S | | ATG3 H236A | |
|---|---|---|---|---|---|---|
| | | | **Set 1** | **Set 2** | **Set 1** | **Set 2** |
| AB Size | Observed AB cross-sectional radii | mean (nm) | 135.3 | 110.3 | 144.9 | 130.8 |
| | | st. dev. (nm) | 46.3 | 32.5 | 43.9 | 40.0 |

| | | | | | | |
|---|---|---|---|---|---|---|
| | Estimated original lognormal distribution of AB sizes | mu | 4.86 | 4.6 | 5.01 | 4.86 |
| | | sigma | 0.324 | 0.33 | 0.260 | 0.286 |
| | | mean original radius (nm) | 136.7 | 105.1 | 155.5 | 133.9 |
| | | st. dev. of orig. radii (nm) | 45.6 | 35.6 | 41.2 | 39.1 |
| | | mean orig. volume (nm$^3$) | $1.5 \times 10^7$ | $6.7 \times 10^6$ | $1.9 \times 10^7$ | $1.3 \times 10^7$ |
| Vacuole Size | Observed vacuole cross-sectional radii | mean (nm) | 859.3 | 767.9 | 938.4 | 815.8 |
| | | st. dev. (nm) | 487.4 | 259.2 | 275.8 | 256.8 |
| | | sample size | 206 | 313 | 193 | 441 |
| | Estimated original lognormal distribution of vacuole sizes | mu | 6.86 | 6.72 | 7.00 | 6.80 |
| | | sigma | 0.291 | 0.294 | 0.209 | 0.261 |
| | | mean original radius (nm) | 999 | 862 | 1177 | 929 |
| AB's per vacuole | Observed AB cross-sections per vacuole | total AB cross-sections | 1627 | 759 | 2553 | 2190 |
| | | vacuoles counted | 203 | 204 | 272 | 201 |
| | | observed ABs / vacuole | 8.01 | 3.72 | 9.39 | 10.9 |
| | Estimated original number of ABs per vacuole | | 52.2 | 29.8 | 59.1 | 66.45 |

**Table S2: Estimations of original AB size and number for Atg3 mutants using the AVS python simulation**

Raw TEM data is the same as used in Table S1, but analyzed using a different estimation method described in Bleta et al. 2026[2]

| | | ATG3 WT | | T213S | | H236A | |
|---|---|---|---|---|---|---|---|
| | | **Set 1** | **Set 2** | **Set 1** | **Set 2** | **Set 1** | **Set 2** |
| AB size | Mu | 5.2 | 5.1 | 4.9 | 4.7 | 5.05 | 4.9 |
| | Sigma | 0.2 | 0.2 | 0.3 | 0.3 | 0.2 | 0.3 |
| | mean original radius (nm) | 184.9 | 167.3 | 140.5 | 115.0 | 159.2 | 140.5 |
| | st. dev. of original radii (nm) | 37.4 | 33.8 | 43.1 | 35.3 | 32.1 | 43.1 |
| | mean original volume (nm$^3$) | $2.9 \times 10^7$ | $2.2 \times 10^7$ | $1.5 \times 10^7$ | $8.3 \times 10^6$ | $1.9 \times 10^7$ | $1.5 \times 10^7$ |
| AB number | Mu | 4.52 | 4.57 | 4.19 | 3.34 | 4.14 | 4.50 |
| | Sigma | 0.4 | 0.4 | 0.4 | 0.4 | 0.4 | 0.4 |
| | Estimated original number AB's per vacuole | 99.5 | 104.5 | 71.5 | 12.7 | 68.0 | 97.5 |

**Table S3: Observed and estimated parameters from Atg10 TEM data**

Estimates based on stereological calculations, as described in Backues et al. 2014[26]

| | | | atg10Δ | | ATG10 WT | |
|---|---|---|---|---|---|---|
| | | | **Set 1** | **Set 2** | **Set 1** | **Set 2** |
| AB Size | Observed AB cross-sectional radii (nm) | mean | N/A | 132.0 | 144.1 | 142.9 |
| | | st. dev. | N/A | 67.8 | 39.5 | 42.6 |
| | Estimated original lognormal distribution of AB sizes | mu | N/A | N/A | 5.04 | 5.00 |
| | | sigma | N/A | N/A | 0.221 | 0.259 |
| | | mean original radius (nm) | N/A | N/A | 157.7 | 153.1 |
| | | st. dev. of original radii (nm) | N/A | N/A | 35.3 | 40.3 |
| | | mean original volume ($nm^3$) | N/A | N/A | $1.9x10^7$ | $1.8x10^7$ |
| Vacuole Size | Observed vacuole cross-sectional radii (nm) | mean | 878.6 | 754.7 | 1061.2 | 946.6 |
| | | st. dev. | 425.6 | 334.8 | 395.5 | 372.7 |
| | | sample size | 239 | 421 | 213 | 420 |
| | Estimated original lognormal distribution of vacuole sizes | mu | 6.74 | 6.66 | 7.10 | 6.87 |
| | | sigma | 0.460 | 0.366 | 0.276 | 0.369 |
| | | mean original radius (nm) | 942 | 836 | 1261 | 1031 |
| ABs per vacuole | Observed AB cross-sections per vacuole | Total AB cross-sections | 6 | 124 | 4165 | 3734 |
| | | vacuoles counted | 203 | 199 | 330 | 311 |
| | | observed ABs / vacuole | 0.03 | 0.62 | 12.6 | 12.0 |
| | Estimated original number of ABs per vacuole | | N/A | N/A | 90.5 | 70.4 |

| | | | ATG10 Single | | ATG10 Double | |
|---|---|---|---|---|---|---|
| | | | **Set 1** | **Set 2** | **Set 1** | **Set 2** |
| AB Size | Observed AB cross-sectional radii (nm) | mean | 148.8 | 144.2 | 110.4 | 107.8 |
| | | st. dev. | 43.9 | 43.3 | 31.2 | 33.9 |
| | Estimated original lognormal distribution of AB sizes | mu | 5.06 | 5.01 | 4.597 | 4.45 |
| | | sigma | 0.245 | 0.260 | 0.317 | 0.374 |
| | | mean original radius (nm) | 161.8 | 154.6 | 104.3 | 92.0 |
| | | st. dev. of original radii (nm) | 40.3 | 40.9 | 33.8 | 35.6 |
| | | mean original volume ($nm^3$) | $2.1x10^7$ | $1.9x10^7$ | $6.4x10^6$ | $5.0x10^6$ |
| Vacuole Size | Observed vacuole cross-sectional radii (nm) | mean | 877.2 | 883.6 | 829.2 | 777.7 |
| | | st. dev. | 406.2 | 336.8 | 386 | 307.1 |
| | | sample size | 196 | 342 | 238 | 366 |
| | Estimated original lognormal distribution of vacuole sizes | mu | 6.78 | 6.80 | 6.71 | 6.63 |
| | | sigma | 0.427 | 0.347 | 0.417 | 0.369 |
| | | mean original radius (nm) | 960 | 956 | 897 | 813 |
| ABs per vacuole | Observed AB cross-sections per vacuole | total AB cross-sections | 2729 | 3142 | 389 | 365 |
| | | vacuoles counted | 311 | 318 | 229 | 201 |
| | | observed ABs / vacuole | 8.77 | 9.88 | 1.70 | 1.81 |
| | Estimated original number of ABs per vacuole | | 46.6 | 52.8 | 15.0 | 17.7 |

**Table S4: Estimations of original AB size and number for Atg10 mutants using the AVS python simulation** Raw TEM data is the same as used in Table S3, but analyzed using a different estimation method described in Bleta et al. 2026[2]

| | | ATG10 WT | | Single | | Double | |
|---|---|---|---|---|---|---|---|
| | | **Set 1** | **Set 2** | **Set 1** | **Set 2** | **Set 1** | **Set 2** |
| AB size | Mu | 5.1 | 5.1 | 5.1 | 5.1 | 4.65 | 4.5 |
| | Sigma | 0.2 | 0.2 | 0.2 | 0.2 | 0.3 | 0.4 |
| | mean original radius (nm) | 167.3 | 167.3 | 167.3 | 167.3 | 109.4 | 97.5 |
| | st. dev. of original radii (nm) | 33.8 | 33.8 | 33.8 | 33.8 | 33.6 | 40.6 |
| | mean original volume ($nm^3$) | $2.2x10^7$ | $2.2x10^7$ | $2.2x10^7$ | $2.2x10^7$ | $7.2x10^6$ | $6.3x10^6$ |
| AB number | Mu | 4.68 | 4.60 | 4.25 | 4.30 | 2.62 | 3.09 |
| | Sigma | 0.4 | 0.4 | 0.4 | 0.4 | 0.4 | 0.4 |
| | Estimated original number AB's per vacuole | 116.7 | 107.8 | 75.9 | 79.8 | 14.4 | 23.8 |

**Table S5: Olinucleotide Primers Used in this Study**

Capital letters indicate portions of the primer designed to anneal to the template; lowercase letters are additions for downstream cloning.

| Number | Name | Sequence |
|---|---|---|
| SKB19 | M13 For | 5'-GTAAAACGACGGCCAG-3' |
| SKB20 | M13 rev | 5'-CAGGAAACAGCTATGAC-3' |
| SKB309 | ADH1 term rev | 5'-ccgggctgcaggaattcGGTGTGGTCAATAAGAGCGAC-3' |
| SKB350 | PEP4 checking F | 5'-AACCAAGTTGCTGCAAAAGTC-3' |
| SKB351 | PEP4D::leu checking R | 5'-TAGTCGACCTCCTCTTCAGTAC-3' |
| SKB352 | PEP4 checking R | 5'-GTATTTACGCAAGAAGGCATCAC-3' |
| SKB398 | Atg10 F2 | 5'-CAGTATTTTCATATTTAGTTGGTTAGGTTATGAAGATTC Acggatccccgggttaattaa-3' |
| SKB399 | Atg10 R1 | 5'-gatgattgcatagtgttttaaaaagctttcctaggttaagGAATTCGAGCT CGTTTAAAC-3' |
| SKB400 | Atg3 F2 | 5'-tacaccgagtatacaacatgactataccatggaaggttggCGGATCCC CGGGTTAATTAA-3' |
| SKB401 | Atg3 R1 | 5'-accacctggcttgcagctaatagtgaaaaaacacaaatttGAATTCGA GCTCGTTTAAAC-3' |
| SKB421 | Atg3 p | 5'-acggtatcgataagcttTTCGGTGGGTCCAGACCTGT-3' |
| SKB422 | Atg3 reverse | 5'-ccccgggctgcaggaattcTTTAAATGATAAAGCTCAGA-3' |
| SKB425 | Atg3 lin For | 5'-AGTACGCCTGCGGGGGGGAC-3' |
| SKB426 | Atg3 lin Rev | 5'-TTGCACATGTTCAGTTTCAGATCCTATGTATTCCAGAA CATC-3' |
| SKB427 | Atg3-FR-PA Forward | 5'-tctgaaactgaacatgtgcaaCTTGCGCAACACGATGAAGCC- 3' |
| SKB429 | Atg3 - 50bp F | 5'-CTTGACGTATATCAAGCTAGC-3' |
| SKB434 | Atg10 F1 | 5'-aaaaaaaaaaaagggctaaaaaacagaattatcagacttgCGGATCC CCGGGTTAATTAA-3' |
| SKB441 | Atg3-FR-PA Reverse (n-term) | 5'-gtcccccccgcaggcgtactGCTTGATATCGAATTCGCGT-3- |
| SKB444 | Pringle Internal Checking R | 5'-TTAATTAACCCGGGGATCCG-3' |
| SKB477 | Atg3 mut H232A F | 5'-GTTTCCATTgcTCCATGTAAGCATGCTAATGTAATGAAA -3' |
| SKB478 | Atg3 mut H232A R | 5'-TACATGGAgcAATGGAAACAGATAACACTGAATTCTTG -3' |
| SKB481 | Atg10 seq +200 bp rev | 5'-GGGCTCGTTATATACTTTTGAATATGTA-3' |
| SKB482 | Atg10 -324 new seq flanking Forward | 5'-GGCGACAAGGTGTTCTAGT-3' |
| SKB488 | Atg10 mut H131A F | 5'-TATTCTTTCgcaCCATGCGATACATCATGTATAGTAGGT GA-3' |
| SKB489 | Atg10 mut H131A R | 5'-TATCGCATGGtgcGAAAGAATACCAAACGCTACCT-3' |
| SKB490 | NEW Atg10 mut Y73Q F | 5'-CAAAAGTAcAaAACGAGCCCTTACTACTTT-3' |

| | | |
|---|---|---|
| SKB491 | NEW Atg10 mut Y73Q R | 5’-GGGCTCGTTtTgTACTTTTGAATATGTAAG-3’ |
| SKB492 | Atg3 mut Y179F F | 5’-CGACATCTtTTAGGGTCCCTAAAATGTATATAGTGGGGT-3’ |
| SKB493 | Atg3 mut Y179F R | 5’-TTAGGGACCCTAAaAGATGTCGAGTACGCAATA-3’ |
| SKB494 | Atg3 mut Y179S F | 5’-CGACATcTTCTAGGGTCCCTAAAATGTATATAGTGGGGT-3’ |
| SKB495 | Atg3 mut Y179S R | 5’-TAGGGACCCTAGAAgATGTCGAGTACGCAAT-3’ |
| SKB547 | ATG5pro cloning F | 5’ataagcttgatatcgaattcATAACTTCTTCAAATACAGACTCTATTATTCAAGAA-3’ |
| SKB548 | Cyc1 term cloning R | 5’-gctctagaactagtggatccGCAAATTAAAGCCTTCGAGCG -3’ |

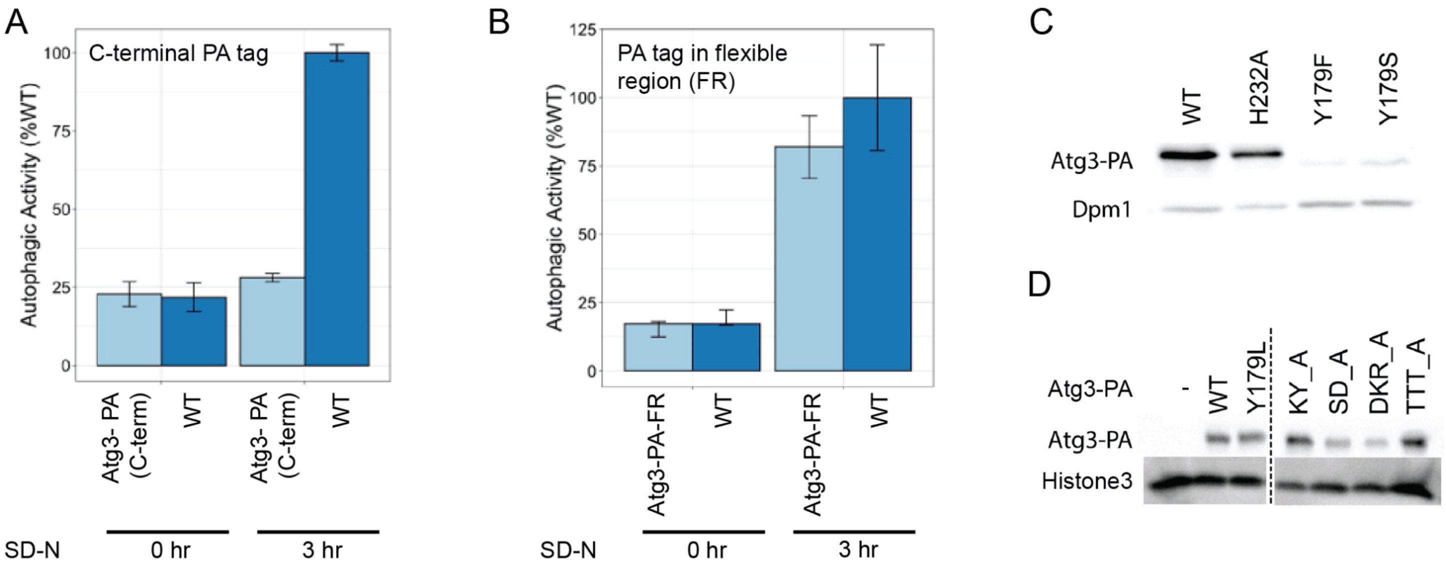


**Figure S1:** Atg3 tagged with Protein A in the flexible region is functional and can be used for monitoring the expression levels of active site mutants. (**A**) Addition of a Protein A (PA) tag to the C-terminus of Atg3 causes a loss of autophagic activity. The pho8Δ60 assay was performed in parallel on a wild type strain and an atg3Δ strain with Atg3-PA (C-terminal tag) with 0 hours and 3 hours of nitrogen starvation to induce bulk autophagy. N=2 biological replicates. (**B**) Insertion of a PA tag within the flexible region (FR) of Atg3, between residues Q117 and S118, allows for retention of autophagic activity. The pho8Δ60 assay was performed in parallel on a wild type strain and an atg3Δ strain with Atg3-PA-FR (PA tag in the FR) with 0 hours and 3 hours of nitrogen starvation to induce bulk autophagy. N=2 biological replicates. Error bars = 95% confidence intervals. (**C and D**) Expression levels of WT and mutant Atg3-PA (PA tag in FR region) in strains used for assessing autophagy in figure 1A-B. The Y179F mutant shows normal activity despite reduced expression levels.

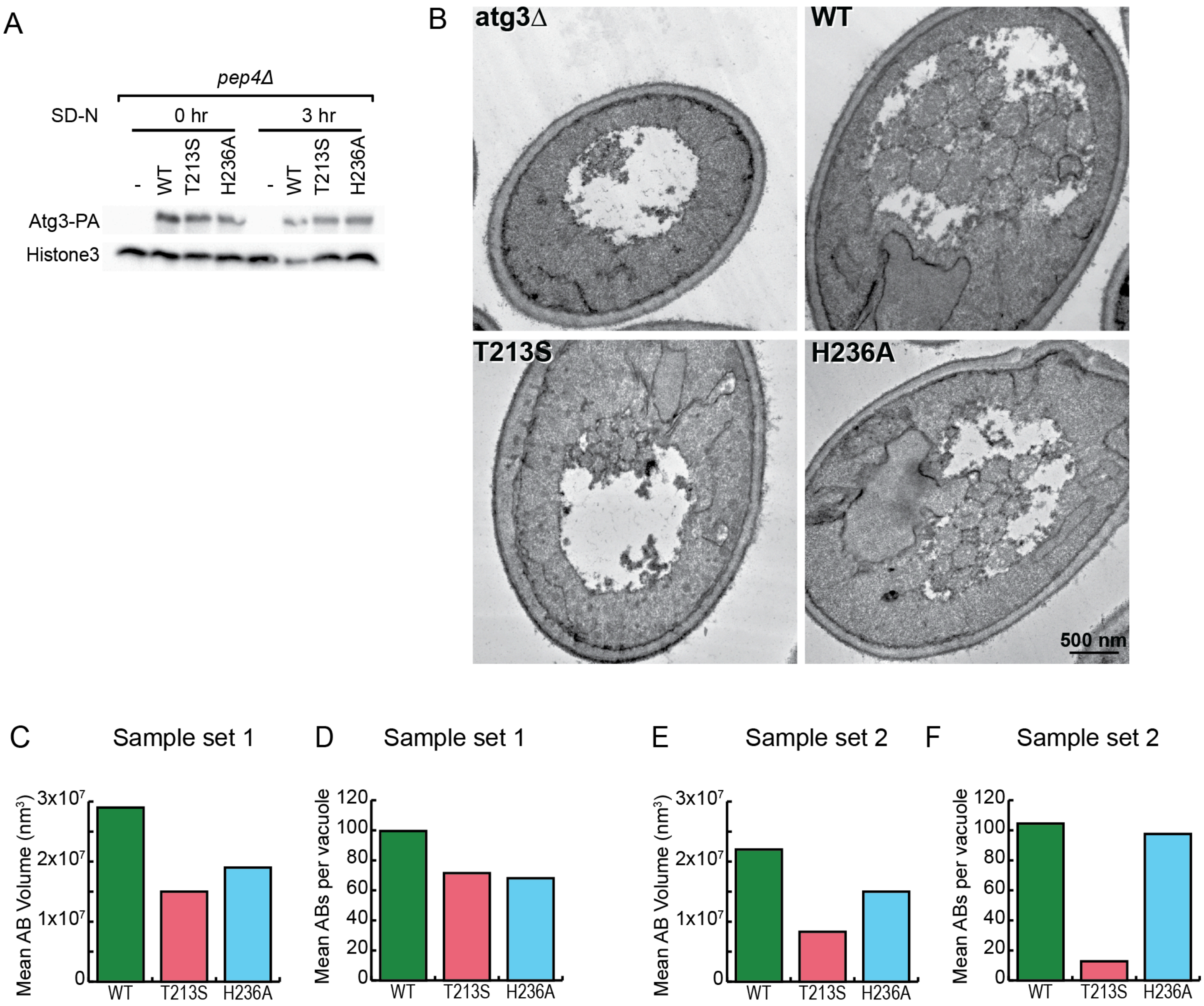


**Figure S2:** A second set of TEM images and an alternate analysis support the conclusion that Atg3 affects body size with a smaller effect on body number. (**A**) Expression levels of WT and mutant Atg3-PA in the actual cultures used for the TEM images shown in panel B. (**B**) Representative TEM images of the second set of samples. This independent biological replicate used the same strains as the first sample set (shown in figure 2 D), but was grown up as a separate set of cultures on a different day. Starvation for 3 hours in SD-N was used to induce autophagy. Data from this sample set is shown in Figure 2 G, H, K and L. (**C-F**) Estimation of the original volume and number of ABs in both TEM sample sets via an alternative simulation-based method supports the conclusion that Atg3 affects both AB size and number, but with a larger effect on AB size - only one mutant (T213S) in one sample set (set 2) shows a larger estimated effect on AB number than size

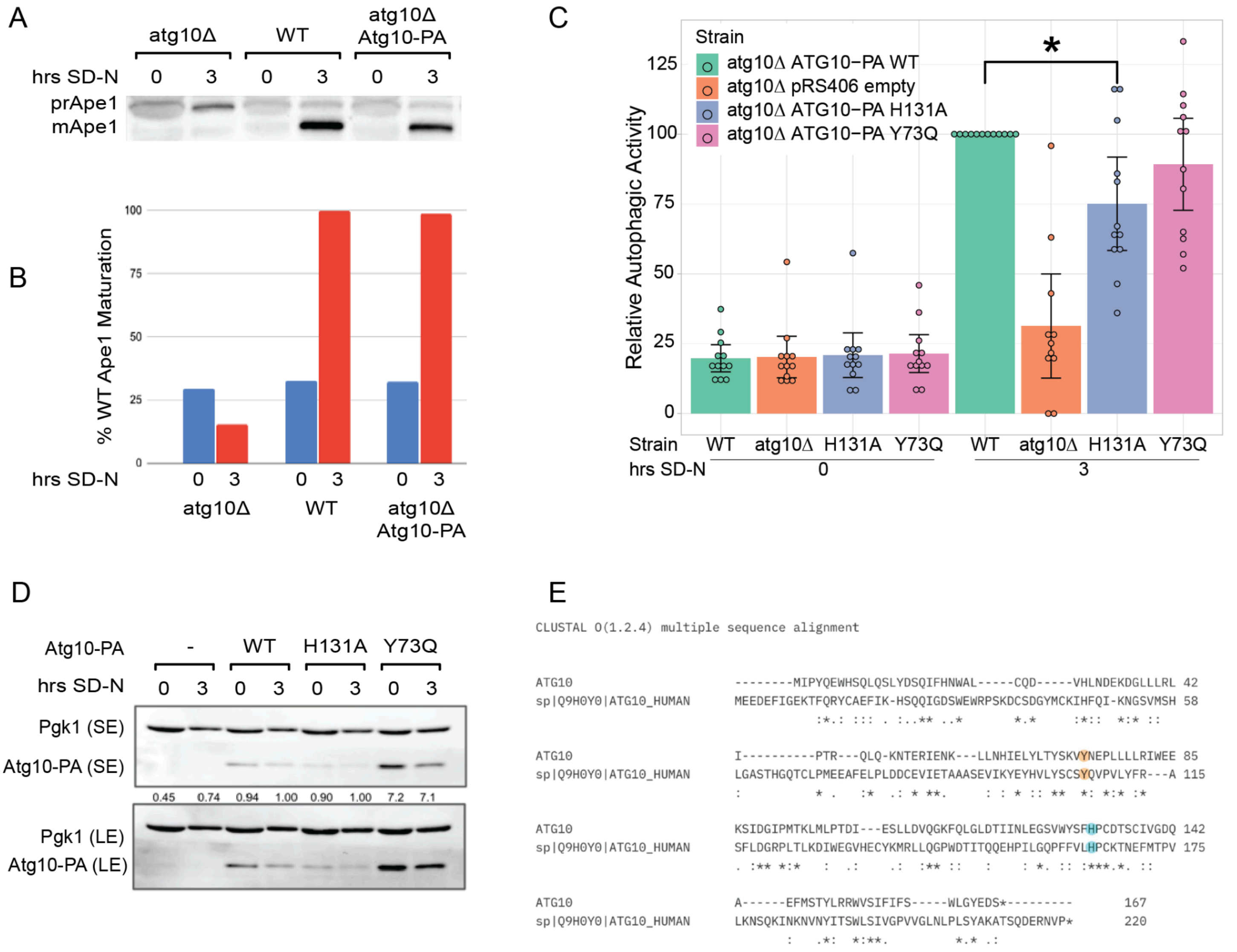


**Figure S3:** C-terminally tagged Atg10 is functional while the Y73Q mutation causes overexpression without a significant effect on autophagic activity. (**A**) Addition of Protein A (PA) tag to the C-terminus of Atg10 does not disrupt Ape1 processing. Ape1 processing was assessed by western blot in a WT strain, an *atg10Δ* strain, and an *atg10Δ* strain expressing Atg10-PA, with 0 hours and 3 hours of nitrogen starvation. (**B**) Quantification of A. Ape1 processing is measured as % of total Ape1 in the mature form, normalized to the WT 3 hr sample. N=1. (**C**) Autophagic activity, as measured by the pho8Δ60 assay, of the Atg10 H131A and Y73Q single mutants. Only H131A is significantly different from the WT. N=12, Error bars = 95% confidence intervals. * = significantly different ($p < 0.05$) from both WT (1-sample T-test) and *atg3Δ* (ANOVA with Tukey HSD). (**D**) Expression levels of WT and mutant Atg10-PA in the strains used in panel C. Y73Q leads to much higher levels of Atg10 protein, while H131A is expressed at levels comparable to the WT. SE = short exposure, LE = long exposure; quantifications are the ratio of Atg10-PA to Pgk1, normalized to the WT 3hr sample. (**E**) Alignment of the amino acid sequence of *S. cerevisiae* Atg10 (top line) with human Atg10 (bottom line), showing conservation of Y73 as human Y106 (orange) and H131 as human H164 (teal).

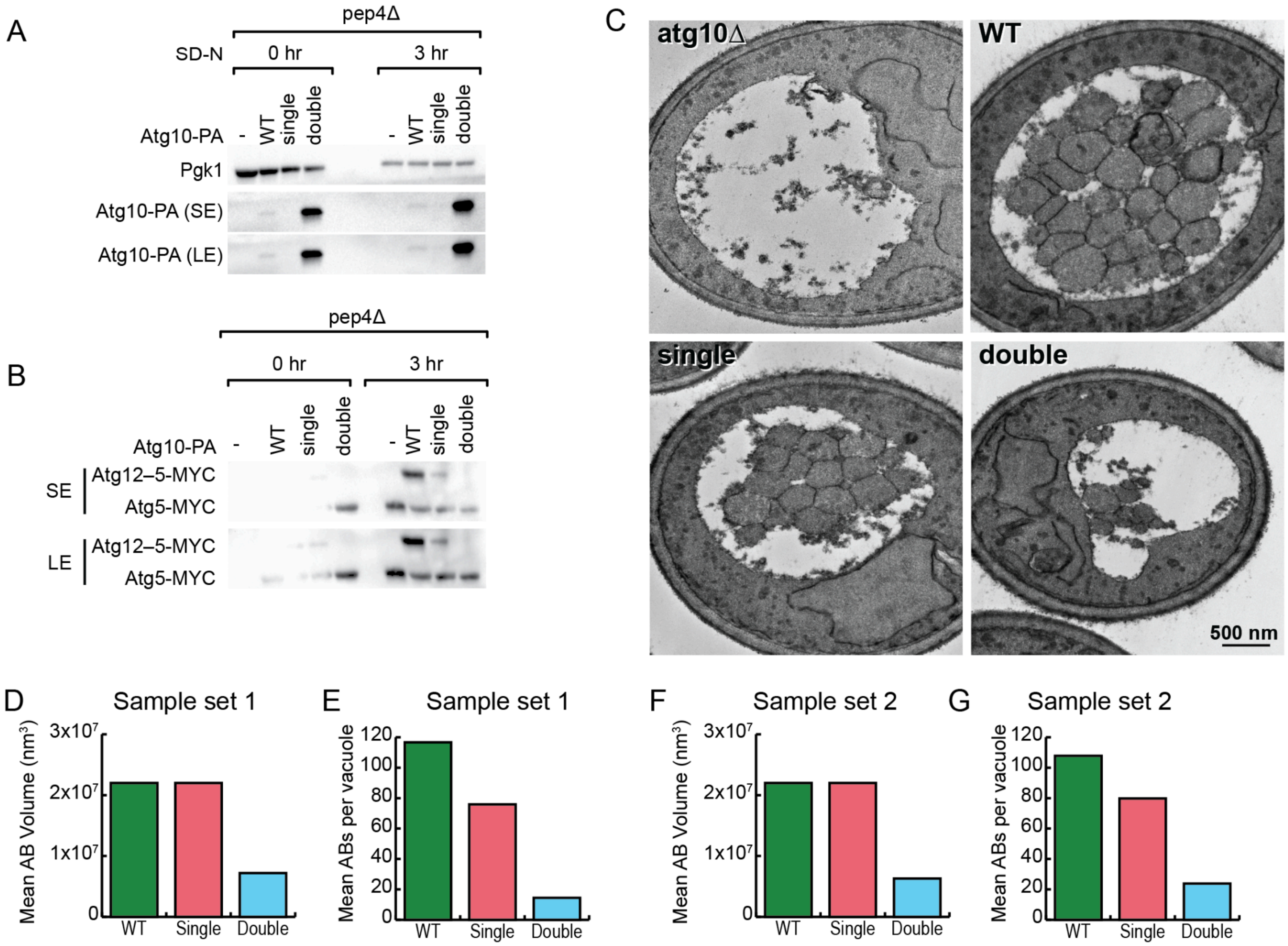


**Figure S4:** A second set of TEM images and an alternate analysis support the conclusion that Atg10 primarily affects body number. (**A and B**) Western blots showing Atg10-PA expression levels and Atg12–5-MYC conjugation in the cultures that generated the TEM images shown Figure 4D (sample set 1), validating the identity of these strains. (**C**) Representative TEM images of the second set of samples. This independent biological replicate used the same strains as the first sample set (shown in fig 4D), but was grown up as a separate set of cultures on a different day. Starvation for 3 hours in SD-N was used to induce autophagy. Data from this sample set is shown in figure 4 I, J, M and N. (**D-G**) Estimation of the original volume and number of ABs in both TEM sample sets via an alternative simulation-based method supports the conclusion that the Atg10 H131A single mutant affects only AB number, not size, while the more severe Y73Q, H131A double mutant reduces both AB size and number.
